# Ribosome collisions in bacteria promote ribosome rescue by triggering mRNA cleavage by SmrB

**DOI:** 10.1101/2021.08.16.456513

**Authors:** Kazuki Saito, Hanna Kratzat, Annabelle Campbell, Robert Buschauer, A. Maxwell Burroughs, Otto Berninghausen, L. Aravind, Roland Beckmann, Rachel Green, Allen R. Buskirk

## Abstract

Ribosome rescue pathways recycle stalled ribosomes and target problematic mRNAs and aborted proteins for degradation. In bacteria, it remains unclear how rescue pathways distinguish ribosomes stalled in the middle of a transcript from actively translating ribosomes. In a genetic screen in *E. coli*, we discovered a novel rescue factor that has endonuclease activity. SmrB cleaves mRNAs upstream of stalled ribosomes, allowing the ribosome rescue factor tmRNA (which acts on truncated mRNAs) to rescue upstream ribosomes. SmrB is recruited by ribosome collisions; cryo-EM structures of collided disomes from *E. coli* and *B. subtilis* reveal a distinct and conserved arrangement of the individual ribosomes and the composite SmrB binding site. These findings reveal the underlying mechanism by which ribosome collisions trigger ribosome rescue in bacteria.

## Introduction

Ribosomes often encounter obstacles that stop them in their tracks: the synthesis of roughly 1 out of every 250 proteins in *E. coli* ends in failure (*1*). Problems arise in many ways. Ribosomes arrest when they reach the 3’-end of transcripts that lack a stop codon, for example, due to premature transcriptional termination or mRNA decay (*2*). Ribosomes also stall on intact messages when they encounter chemical damage in the mRNA (*2, 3*), codons that are decoded very slowly (*4*), or specific nascent peptide sequences that inhibit their own translation (*5, 6*). Although stalling is sometimes resolved productively, prolonged pauses often lead to aborted protein synthesis. Such translational failures are dangerous because they trap ribosomes in inactive complexes and produce incomplete proteins that can be toxic. To meet these challenges, bacteria have evolved ribosome rescue mechanisms that selectively recognize stalled ribosomes, rescue the ribosomal subunits, and target problematic mRNAs and nascent proteins for degradation (reviewed in (*7*)).

How do ribosome rescue factors in bacteria distinguish stalled ribosomes from actively translating ribosomes? There are two rescue pathways active in *E. coli*: the main one (mediated by tmRNA and its protein partner, SmpB) and a backup pathway (mediated by ArfA) that is induced when tmRNA is overwhelmed (*8*). These pathways differ in fundamental ways: the tmRNA-SmpB complex enters ribosomes and encodes a short peptide tag to target the nascent peptide for proteolysis (*9*) whereas ArfA simply promotes peptidyl hydrolysis by recruiting the canonical release factor RF2 (*10*). Because both factors bind in the mRNA channel, their activity is inhibited by intact mRNA downstream of the stall site (*7, 11–13*). In cases where translation stalls on intact messages, the current model is that mRNA cleavage yields truncated mRNAs that become good substrates for rescue pathways. For example, ribosome stalling during termination in an inefficient context, Glu-Pro-stop (EP*), leads to mRNA cleavage in the ribosomal A site (*14*). mRNA cleavage is an attractive mechanism that could explain how rescue factors identify ribosomes that have irreversibly arrested. Yet the signal that triggers mRNA cleavage and the identity of the nucleases involved have remained elusive.

Recent work has revealed insights into how eukaryotic cells recognize and rescue stalled ribosomes. The E3 ubiquitin ligase Hel2 was identified in a genetic screen in yeast as a factor that promotes ribosome rescue by adding ubiquitin to specific ribosomal proteins (*15*). Hel2 is thought to recognize the interface between two small ribosomal subunits formed when an upstream ribosome collides into a stalled ribosome (*16, 17*). Collisions are the trigger that leads to ubiquitination of r-proteins and activation of downstream quality control pathways including subunit splitting, mRNA decay, and degradation of the nascent polypeptide (*18*). In the absence of ubiquitination systems and Hel2 homologs, however, it has been unclear whether similar mechanisms are at play in bacteria.

Here, we report that the *E. coli* protein SmrB cleaves mRNA upstream of stalled ribosomes, promoting ribosome rescue. SmrB contains an SMR domain associated with nuclease activity (*19–21*); SMR-domain proteins are broadly conserved in bacteria and eukaryotes as well as in a few archaeal lineages, one of the few rescue factors with a widespread presence across the three superkingdoms of life. We show that following SmrB cleavage at the 5’-boundary of the stalled ribosome, upstream ribosomes translate to the 3’-end of the upstream mRNA fragment and are rapidly rescued by tmRNA. We show that SmrB binds preferentially to collided ribosomes and only cleaves at stalling motifs where collisions occur, arguing that SmrB recognizes aberrant translation through recruitment to collided ribosomes. Moreover, we present cryo-EM structures of ribosome dimers stalled at specific stalling motifs in *B. subtilis* and *E. coli*, revealing a distinct and conserved architecture of disomes formed by collisions. In SmrB-bound disomes we define a composite binding site that explains how SmrB is specifically recruited to and activated by ribosome collisions. These findings establish that ribosome collisions are the trigger for ribosome rescue in *E. coli* through recruitment of the nuclease SmrB.

## Results

### A genetic selection for novel rescue factors

To identify novel factors that act early in the ribosome rescue pathway in *E. coli*, we performed a genetic selection similar to the one used previously to discover Hel2 in eukaryotes (*15*), searching for mutants that allow ribosomes to translate through a strong stalling motif and complete the translation of a downstream ORF. Our selection is based on a reporter construct encoding a fusion of NanoLuc upstream of the bleomycin resistance protein (Ble) that will confer growth on selective media (Fig 1A). Two control constructs are shown: one has a stop codon in between the genes and produces NanoLuc alone (Stop); the second is a direct fusion without any intervening stalling motif (Non-stall) and produces full-length fusion protein. In a third construct, we inserted the strong SecM arrest motif at the junction of NanoLuc and Ble (SecM); overexpression of reporters containing this stalling motif leads to mRNA cleavage and tagging of the nascent peptide by tmRNA (*5*).

**Figure 1.**
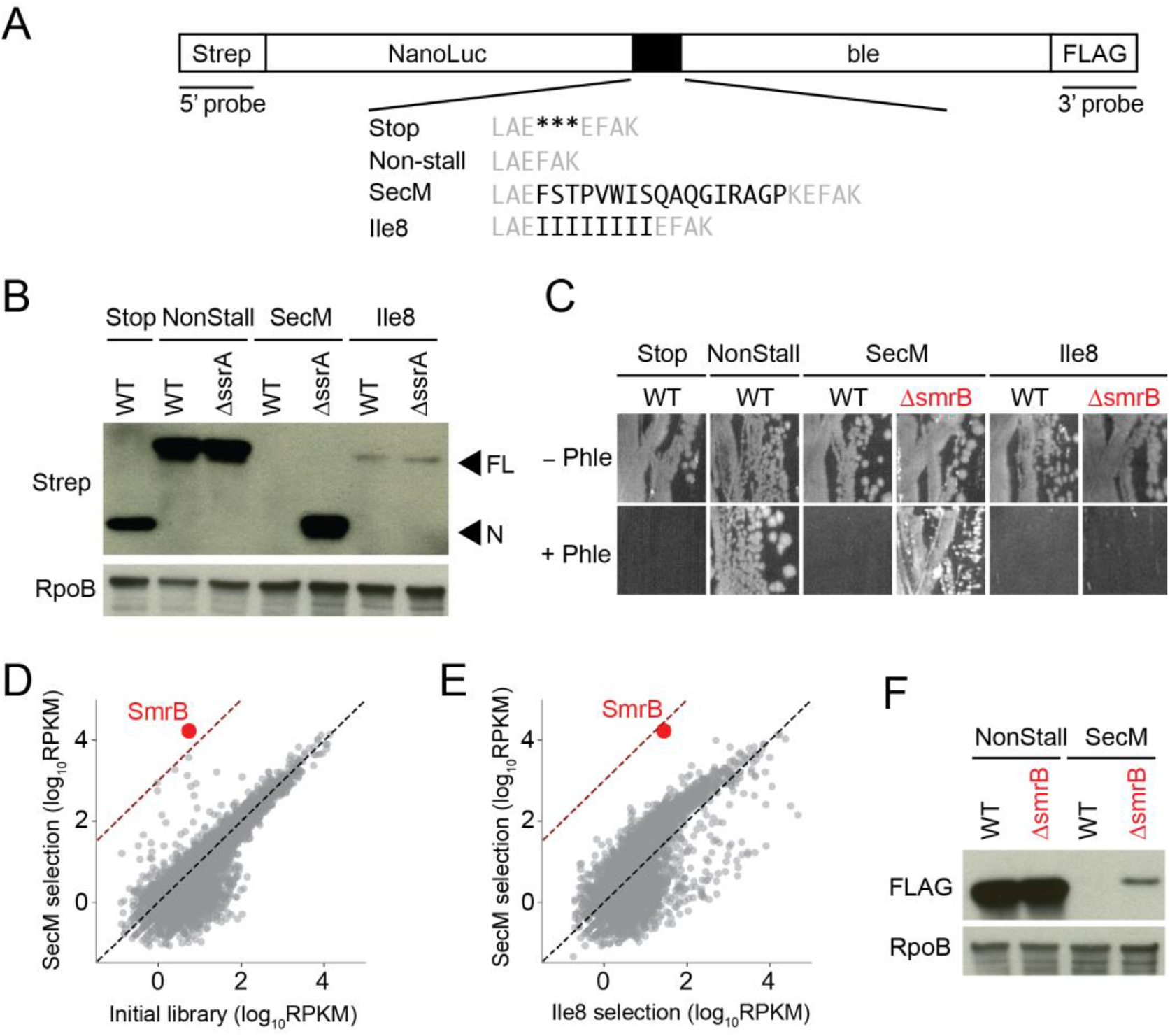
A genetic selection for novel rescue factors. (**A**) Reporters for studying ribosome rescue in *E. coli*. Between the NanoLuc gene and bleomycin resistance gene, we inserted either stop codons, no added sequence, the SecM stalling motif, or eight Ile codons (black). (**B**) Reporter protein from wild-type and ΔssrA strains was detected by antibodies against the N-terminal Strep-tag. Arrows indicate the full-length fusion protein (FL) and shorter NanoLuc protein (N). The RpoB protein serves as a loading control. (**C**) Growth of wild-type and ΔsmrB strains expressing various reporters on media with or without 50 μg/mL phleomycin. (**D**) and (**E**) The results of TN-seq are shown as a scatter plot of mapped reads (rpkm) for each gene corresponding to the number of transposon insertions. The dashed line indicates 1000-fold enrichment. (**F**) Full-length reporter protein was detected using antibodies against the FLAG-tag.

We first confirmed that the SecM reporter undergoes stalling and ribosome rescue using antibodies against the Strep-tag at the N-terminus of the NanoLuc-Ble fusion protein. Although abundant full-length protein is visible in the Non-stall control, no full-length protein is detectable for the SecM reporter due to the strong arrest sequence (Fig 1B). To visualize the aborted protein product, we introduced the SecM reporter into cells lacking tmRNA (encoded by *ssrA*). Using the anti-Strep antibody, we observed truncated protein that is roughly the same size as NanoLuc produced from the Stop control (Fig 1B).

As another control, we developed a reporter with eight consecutive isoleucine residues at the fusion site; this reporter yields relatively little full-length protein detected by the anti-Strep antibody, likely because the protein is misfolded and degraded (Ile8, Fig 1B). No truncated protein from the Ile8 reporter was observed in the ΔssrA strain. The relatively low levels of full-length protein independent of ribosome stalling and rescue makes the Ile8 reporter a useful control in our genetic selection, as discussed below.

Using the NanoLuc-Ble reporters, we found that expression of the full-length reporter protein from the Non-stall construct conveys resistance to 50 μg/mL phleomycin (an antibiotic structurally related to bleomycin) (Fig 1C). In contrast, cells expressing NanoLuc alone from the Stop construct are sensitive to this concentration of phleomycin, as expected. Next, we tested the SecM and Ile8 constructs and observed that wild-type cells expressing these reporters are also phleomycin sensitive, in line with the finding that the full-length protein is expressed at very low levels from these constructs. These results establish parameters for the screen to identify gene deletions that increase the level of full-length reporter protein.

To perform the genetic selection we adopted a Tn-seq approach (*22, 23*), creating a knock-out library of about 5 million colonies through random insertion of Tn5 transposase into *E. coli* K12 MG1655. We transformed this library with a plasmid expressing the SecM or the Ile8 reporter and plated the transformants on media containing 50 μg/mL phleomycin. After harvesting phleomycin-resistant cells, we counted the number of transposon insertions per gene (normalized by length) in units of reads per kilobase per million mapped reads (rpkm) (Fig 1D). Compared to the initial library, 29 genes in the SecM reporter strain and 109 genes in the Ile8 reporter strain exhibited a more than 10-fold enrichment in transposon insertions. Many of these genes are false positives relevant to phleomycin toxicity. By comparing the results of the SecM selection with the Ile8 selection, we can remove false positives from consideration, focusing instead on genes that are selectively enriched in the SecM selection because they affect ribosome stalling and rescue (Fig 1E).

We found a single gene, *smrB*, where transposon insertions were strongly enriched in the SecM reporter strain compared to the Ile8 reporter strain (~600-fold, Fig 1E). To confirm this phenotype, we constructed a clean *smrB* deletion strain and found that ΔsmrB cells expressing the SecM reporter are resistant to phleomycin whereas cells expressing the Ile8 reporter remain sensitive (Fig 1C). Furthermore, using the anti-FLAG antibody, we observe that deletion of *smrB* yields increased levels of full-length NanoLuc-Ble protein from the SecM reporter (Fig 1F) and from similar reporters with other stalling motifs, such as 12 rare Arg codons or the Glu-Pro-stop motif (EP*) (Fig S1). In contrast, loss of *smrB* has no effect on expression of the Non-stall reporter.

### SmrB is a conserved nuclease involved in ribosome rescue

The *E. coli* SmrB protein is 183 amino acids long and contains a domain of the <u>S</u>mall <u>M</u>utS <u>R</u>elated (SMR) superfamily first proposed to act as a DNase in MutS-like proteins in bacteria and plants. However, more recent studies have shown that SMR domains possess endonucleolytic RNase activity (*19, 24, 25*). Indeed, we recently identified an SMR protein, Cue2, as the endonuclease that cleaves mRNA upstream of stalled ribosomes in yeast (*20*). A systematic analysis of the SMR domains across the tree of life revealed independent fusions to a diverse array of RNA-binding domains supporting the hypothesis that it primarily operates on RNA (*21*).

To better understand the evolutionary trajectories of the SMR proteins, we performed phyletic pattern (Fig 2A) and phylogenetic analyses (Fig S2). We find that SMR domains are broadly conserved in bacteria, though notably underrepresented in the PVC group and the actinobacteria (Fig 2A). They are found across all sampled eukaryotic lineages, but are relatively uncommon in Archaea, where they are found mainly in Asgardarchaeota and Thermoplasmatota. These data suggest that the SMR domain might have entered the eukaryotic stem lineage at some point from a bacterial source.

**Figure 2.**
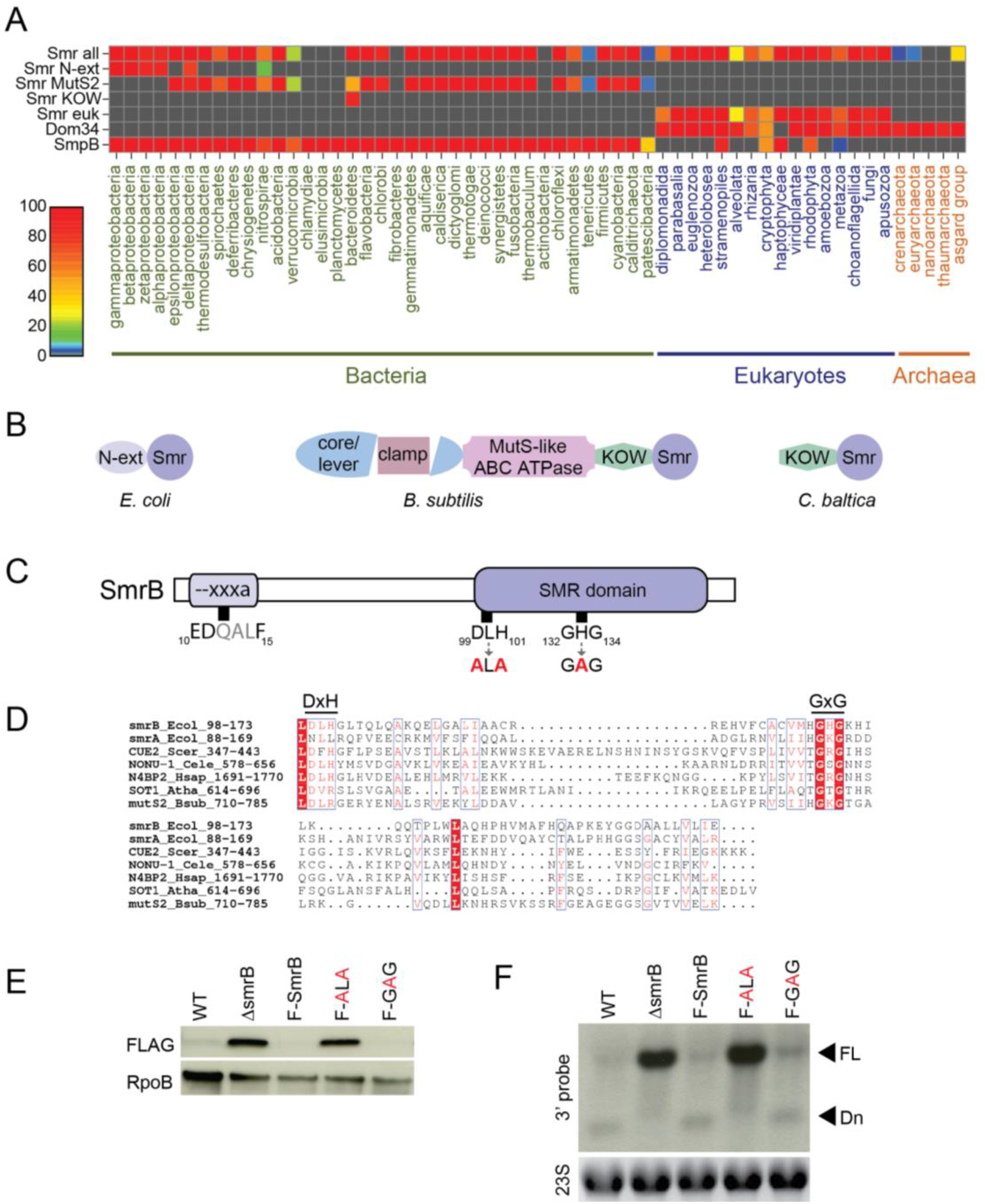
SMR-domain proteins are conserved RNA nucleases. (**A**) Heat map demonstrating the conservation and distribution of SMR-domain proteins and other related translational quality control factors. Smr-all includes all types of SMR-domain proteins; Smr-euk includes only the eukaryotic branch. (**B**) Domain organization of three representative bacterial proteins containing an SMR domain. (**C**) Domain organization of *E. coli* SmrB. Mutations at the two conserved DxH and GxG motifs are indicated by red bold letters. (**D**) Sequence alignment of SMR domains of representative proteins. Identical residues are shown in white with a red background; conserved residues are shown in red. The identity of each sequence is represented by the gene name, species name, and numbers indicating the beginning and the end of the residues used for the alignment. Ecol, *Escherichia coli*; Scer, *Saccharomyces cerevisiae*; Cele, *Caenorhabditis elegans;* Hsap, *Homo sapiens*; Atha, *Arabidopsis thaliana*; Bsub, *Bacillus subtilis*. (**E**) Full-length SecM reporter protein was detected with antibodies against the FLAG-tag. The RpoB protein serves as a loading control. F-SmrB, F-ALA, and F-GAG indicate endogenously FLAG-tagged SmrB and its variants. (**F**) The SecM reporter mRNA was detected on northern blots using a probe which anneals to the 3’-end of the reporter. Ethidium bromide staining of 23S rRNA serves as a loading control. FL = full-length and Dn = the downstream mRNA fragment.

The bacterial SMR domains cluster into three major clades. *E. coli* SmrB together with other proteobacterial versions form the first of these clades (Fig 2B) typified by a characteristic extension N-terminal to the SMR domain. This extension contains a predicted helix (residues 9-20) with a strongly conserved “--xxxa” motif (two negatively charged residues followed by three variable and one aromatic residue; E_10_DQALF_15_ in *E. coli* SmrB; Fig 2C and Fig S3) followed by a poorly conserved, largely unstructured segment. In gammaproteobacteria, like *E. coli*, a duplication led to two copies of these SMR proteins per genome; one with active site residues conserved (SmrB) and the other predicted to be enzymatically inactive (SmrA). The second clade contains the most common bacterial version typified by the *B. subtilis* MutS2 protein. From N- to C-terminus, these proteins contain the core/lever and clamp domains, the MutS DNA mismatch repair protein-type P-loop ABC ATPase domain, as well as an additional KOW domain with a SH3-like fold and a C-terminal SMR domain (Fig 2B). Notably, the MutS2 proteins lack the mismatch recognition and connector domains typical of the canonical MutS protein involved in mismatch repair. We identified several independent occasions where the predicted active site residues have been lost in this clade. The third clade is the smallest, restricted to the *Bacteroidetes* lineage, with an N-terminal KOW domain and C-terminal SMR domain.

An alignment of the SMR domain (residues 98-173) reveals that SmrB contains the conserved residues associated with endonuclease and RNA-binding activity (*21, 26*): residues D_99_LH_101_ correspond to the DxH motif implicated in catalysis and residues G_132_HG_134_ with the GxG motif marking the loop predicted to interact with RNA substrates (Fig 2D). We generated *E. coli* strains where the DxH and GxG motifs were mutated to ALA and GAG, respectively (Fig 2C), at the endogenous *smrB* locus tagged with an N-terminal FLAG epitope. The FLAG tag does not inhibit SmrB activity; like the wild-type, little or no full-length reporter protein is detectable in this strain (Fig 2E). Importantly, we observe that the ALA mutation increased full-length protein to a similar extent as deletion of *smrB*, whereas the GAG mutation had no discernable effect in vivo activity. These results suggest that the DxH motif is critical for SmrB activity while the central residue in the GxG motif is not required, consistent with its lower conservation. We note that loss of a second SMR-domain protein encoded in the *E. coli* genome, SmrA, which lacks the DxH motif did not affect expression of the stalling reporter (Fig S4).

To observe more directly the activity of SmrB on the reporter mRNA in vivo, we performed northern blots with a probe binding to the 3’-end of the reporter construct (Fig 2F). In the wild-type strain, the full-length reporter mRNA is barely detectable while the strongest signal comes from a shorter mRNA fragment whose size suggests that cleavage is occurring somewhere near the SecM motif. In the ΔsmrB strain, the downstream fragment disappears and the levels of the full-length mRNA are dramatically higher. As expected, the strains with FLAG-tagged SmrB and the GAG mutant show robust levels of RNA cleavage, whereas the ALA mutant strain shows high levels of reporter mRNA. These results confirm that SmrB cleaves the reporter mRNA in vivo and reveal that SmrB cleavage is the dominant pathway that targets the reporter mRNA for degradation.

### Reporter mRNAs decayed via multiple pathways

We next asked whether we could detect the upstream fragment using a probe against the 5’-end of the reporter mRNA. We began by comparing the mRNA levels of the SecM-short reporter and the EP* reporter in the wild-type strain versus strains lacking either tmRNA or SmrB or both. Whereas we see little reporter RNA in the wild-type strain (since it is cleaved and degraded), the 5’-probe reveals the upstream fragment from both reporters in the ΔssrA strain (Fig 3A), consistent with prior reports that the loss of tmRNA stabilizes the upstream fragment (*5, 14*). As expected, there is no detectable upstream fragment in the ΔsmrB strain. Surprisingly, however, the upstream fragment is present for both reporters in the ΔssrA ΔsmrB strain (ΔΔ); this unexpected result reveals that this truncated mRNA can be produced by one or more mechanisms that are independent of SmrB. It is unlikely that other endonucleases are responsible, given that the downstream fragment detected by the 3’-probe disappears in strains lacking SmrB (Fig 3A). We speculate that in the absence of SmrB, the upstream fragment is generated by exonucleolytic decay of the mRNA back to the stalled ribosome; *E. coli* has three processive 3’-5’ exonucleases implicated in mRNA decay but lacks 5’-3’ exonucleases (*27*).

**Figure 3.**
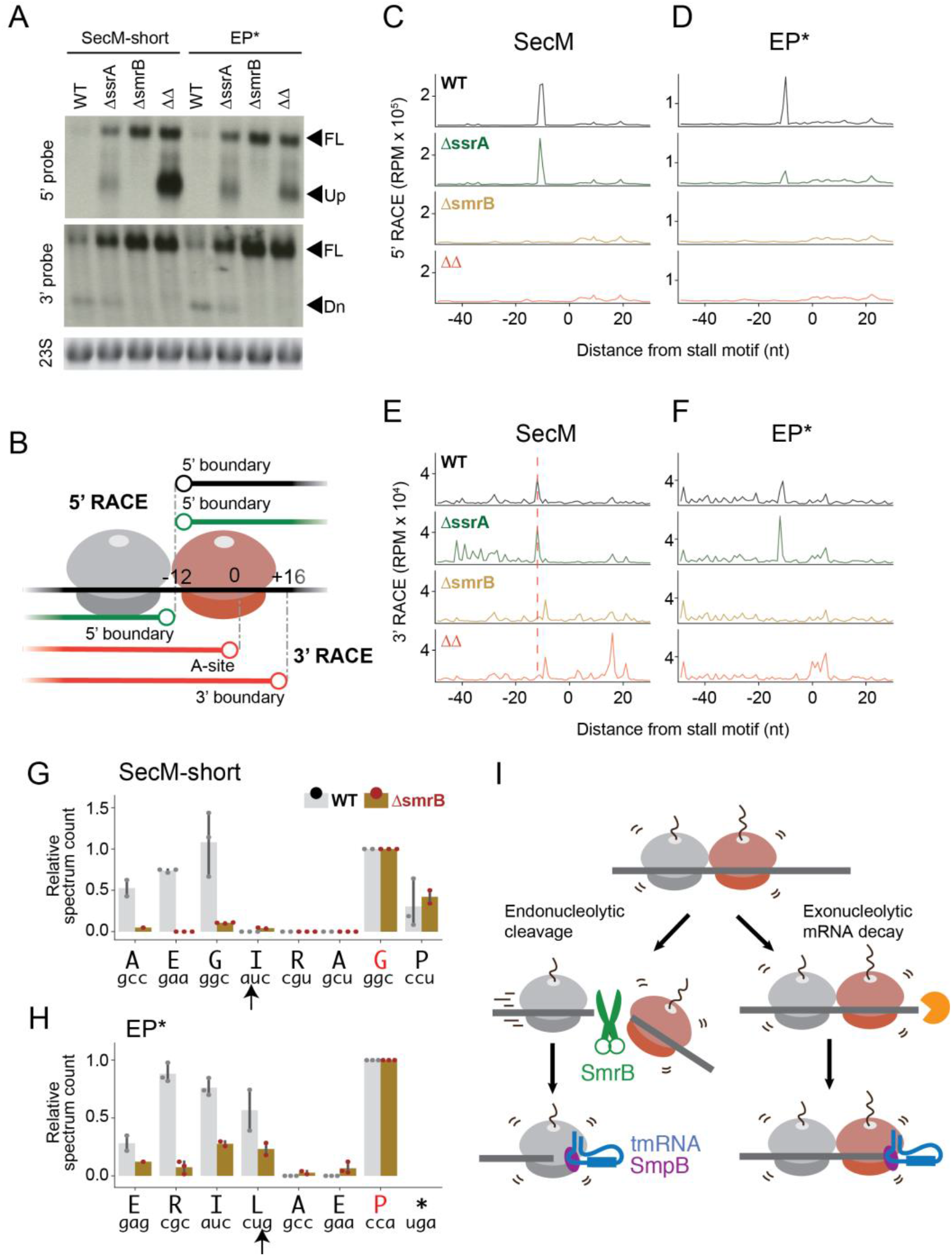
SmrB cleaves at the 5’ boundary of stalled ribosomes. (**A**) Northern blots of reporter mRNA using the 5’-probe and the 3’-probe. Arrows indicate the full-length (FL) or truncated RNAs (upstream or downstream fragments). Ethidium bromide staining of 23S rRNA serves as a loading control. (**B**) Schematic representation of RNA fragments observed in the RACE data. (**C**) and (**D**) The results of 5’ RACE showing the 5’-ends of downstream fragments in reads per million on the reporter sequence. The first nt in the A site codon in the stall motif is designated as zero. (**E**) and (**F**) The results of 3’ RACE showing the 3’-ends of upstream fragments in reads per million on the reporter sequence. The first nt in the A site codon in the stall motif is designated as zero. (**G**) and (**H**) tmRNA tagging near the stall motif in wild-type and ΔsmrB strains. The tagging site is the residue immediately preceding the tmRNA tag in peptide sequences detected by targeted LC-MS-MS. The red letter indicates the residue encoded by the P site codon at the stall site. The arrow indicates the SmrB cleavage site demonstrated by 5’ RACE. The relative spectrum count is shown, normalized by the count at the stall site where tmRNA tagging was expected to occur in both the wild-type and ΔsmrB strains. (**I**) A model for mRNA processing during ribosome rescue. In the endonucleolytic cleavage pathway, SmrB cleaves the mRNA at the 5’ boundary of stalled ribosomes, at the interface of ribosome collisions. Upstream ribosomes then resume translation, reaching the 3’-end of the cleaved mRNA, and are quickly rescued by tmRNA-SmpB. In the secondary pathway, 3’-to-5’ exonucleases degrade mRNA until they hit the 3’ boundary of the stalled ribosome, after which the stalled ribosome is eventually rescued by tmRNA-SmpB.

### SmrB cleaves mRNA at the 5’-boundary of stalled ribosomes

The reporter mRNA is degraded by at least two mechanisms, one that depends on SmrB and another that likely involves exonucleases. To better characterize these pathways, we used RACE to identify the 5’- and 3’-ends of the mRNA fragments produced by these decay events coupled to ribosome stalling (Fig 3B). 5’-RACE experiments reveal the 5’-end of the downstream fragment which is generated solely by endonucleolytic cleavage by SmrB: there is a sharp peak 11 nt upstream of the SecM stall site (Fig 3C). This peak is also evident in the ΔssrA strain, but disappears in the ΔsmrB and the ΔsmrB ΔssrA knockout strains. These data are consistent with the northern blots showing that the downstream fragment is not detectable in strains lacking SmrB (Fig 3A, 3’-probe). We see very similar results from the EP* reporter where the 5’-end of the downstream fragment is 10 nt upstream of the stop codon in the A site of the ribosome (Fig 3D). Based on ribosome footprinting experiments (*28*), we know that the 5’-boundary of the ribosome is roughly 12 nt upstream of the A-site codon, suggesting that SmrB cleaves mRNA at the site where it exits the ribosome.

We also used 3’-RACE to determine the 3’-end of the upstream mRNA fragment (Fig 3B). The data from these experiments are more complex because multiple pathways generate truncated mRNAs near the stall site. For the SecM reporter, in the wild-type and ΔssrA strains, the strongest peak in the 3’-RACE data is 11 nt upstream of the stall site (Fig 3E); in the absence of SmrB, this peak disappears. This same phenomenon was also observed in the EP* reporter (Fig 3F). For both reporters, these positions correspond perfectly with the site of cleavage identified on the downstream fragment by 5’-RACE, suggesting that these upstream fragments are derived from SmrB cleavage at the 5’ boundary of stalled ribosomes.

In strains lacking both SmrB and tmRNA (ΔΔ) the 3’-RACE data provide additional information about other pathways that act on the upstream mRNA fragment. In the absence of both factors, the strongest 3’-RACE signal for the SecM reporter is 16 nt downstream of the first nt in the A site codon, roughly corresponding to the 3’-boundary of the ribosome stalled at the SecM motif (Fig 3E), likely the products of exonucleolytic decay. In contrast, the strongest 3’-RACE signal from the double knockout strain expressing the EP* reporter is at the A-site codon (Fig 3F). These results are broadly consistent with previous reports of mRNA cleavage at both the 5’- and 3’-boundaries of ribosomes stalled on SecM (*5*) and of A-site cleavage within ribosomes stalled during termination at EP* (*14*). Taken together, the 5’- and 3’-RACE data on these stalling reporter mRNAs reveal that SmrB cleaves at the 5’-boundary of stalled ribosomes and that in the absence of SmrB, other pathways lead to mRNA decay up to the A-site codon or the 3’-boundary of the lead stalled ribosome.

### SmrB cleavage liberates stacked ribosomes upstream of stalled ribosomes

Our analyses of the reporter mRNA indicate that SmrB cleaves upstream of stalled ribosomes forming an upstream fragment whose decay is promoted by tmRNA. To ask which ribosome complexes are rescued by tmRNA, we determined where the tmRNA tag is added on both the short-SecM and EP* reporters. We immunoprecipitated reporter protein (using the N-terminal Strep-tag) from both the wild-type and ΔsmrB strains containing a tmRNA variant that tags the reporter protein with a ClpXP-resistant AANDENYAL**DD** sequence (*4*).

We determined the sites of tmRNA tagging by digesting the immunoprecipitated reporter protein with lysyl endopeptidase and subjecting the resulting peptides to LC-MS-MS. For the SecM reporter, where ribosomes stall with the second Gly codon in GIRAGP in the P site (*29*), there is strong signal from the peptide produced when the tmRNA tag is added at the second Gly residue (Fig 3G), as previously reported (*5*). Importantly, robust tagging at this site is observed in both the wild-type and ΔsmrB strains, suggesting that SmrB cleavage is not essential for tagging at the SecM stall site. We also observe strong tagging four residues upstream (at the first Gly) in the wild-type strain (Fig 3G). This result correlates precisely with the SmrB cleavage site determined by RACE; tagging at this site is dramatically reduced in the ΔsmrB strain. Likewise, consistent with previous studies (*6*), we observe tmRNA tagging of the EP* reporter protein at the C-terminal Pro residue in both the wild-type and ΔsmrB strains (Fig 3H). We also observe tagging at the residues upstream of the stalling site at positions where mRNA is cleaved by SmrB in the wild-type strain; this signal is diminished in the ΔsmrB strain.

Taken together, the RACE and tmRNA tagging results lead us to propose the following model for ribosome rescue within ORFs in *E. coli* (Fig 3I). Ribosome stalling leads to endonucleolytic cleavage by SmrB at the 5’-boundary of the first, stalled ribosome (red). No longer impeded by the ribosome trapped on the stalling motif, upstream ribosomes (grey) translate to the end of the upstream mRNA fragment and arrest at the 3’-end generated by SmrB cleavage. The tmRNA-SmpB complex then rescues and releases these ribosomes, consistent with its well-characterized preference for truncated mRNAs. This model highlights how SmrB cleavage and tmRNA activity rapidly clear upstream ribosomes from the message and target it for decay. Consistent with previous studies, tmRNA also releases the initial stalled ribosome (red), tagging the nascent peptide right at the stall site. Decay of the reporter mRNA back to the 3’-boundary of the ribosome (in the case of SecM) or the A-site codon (in the case of EP*) by exonucleases likely allows tmRNA to gain access to these stalled ribosomes (*5, 14, 30*).

### SmrB preferentially binds collided ribosomes

We next asked how SmrB selectively recognizes stalled ribosomes. Although ribosome collisions have not been implicated in bacterial ribosome rescue, our data suggest that SmrB, like Cue2, recognizes collided ribosomes. First, as shown above, 5’-RACE of the downstream fragment in the SecM reporter reveals that SmrB cleavage occurs precisely at the 5’-boundary of the SecM-stalled ribosome; additional peaks are seen (with lower intensity) further upstream of the stall site that cluster in sets centered roughly 25 nt apart, the length of a ribosome footprint (Fig 4A). Second, ribosome profiling data from the ΔssrA ΔsmrB strain expressing a related reporter reveal a strong peak of ribosome density at the SecM stall site, as expected, as well as sets of peaks roughly 25 nt apart extending upstream (Fig 4A). The 5’-boundaries of the SecM stalled ribosome and the first three stacked ribosomes align well with the sites of SmrB cleavage seen in the RACE data. These findings suggest that SmrB cleavage happens in the context of ribosome collisions, arguing that collisions may serve as a signal for SmrB recruitment or activation.

**Figure 4.**
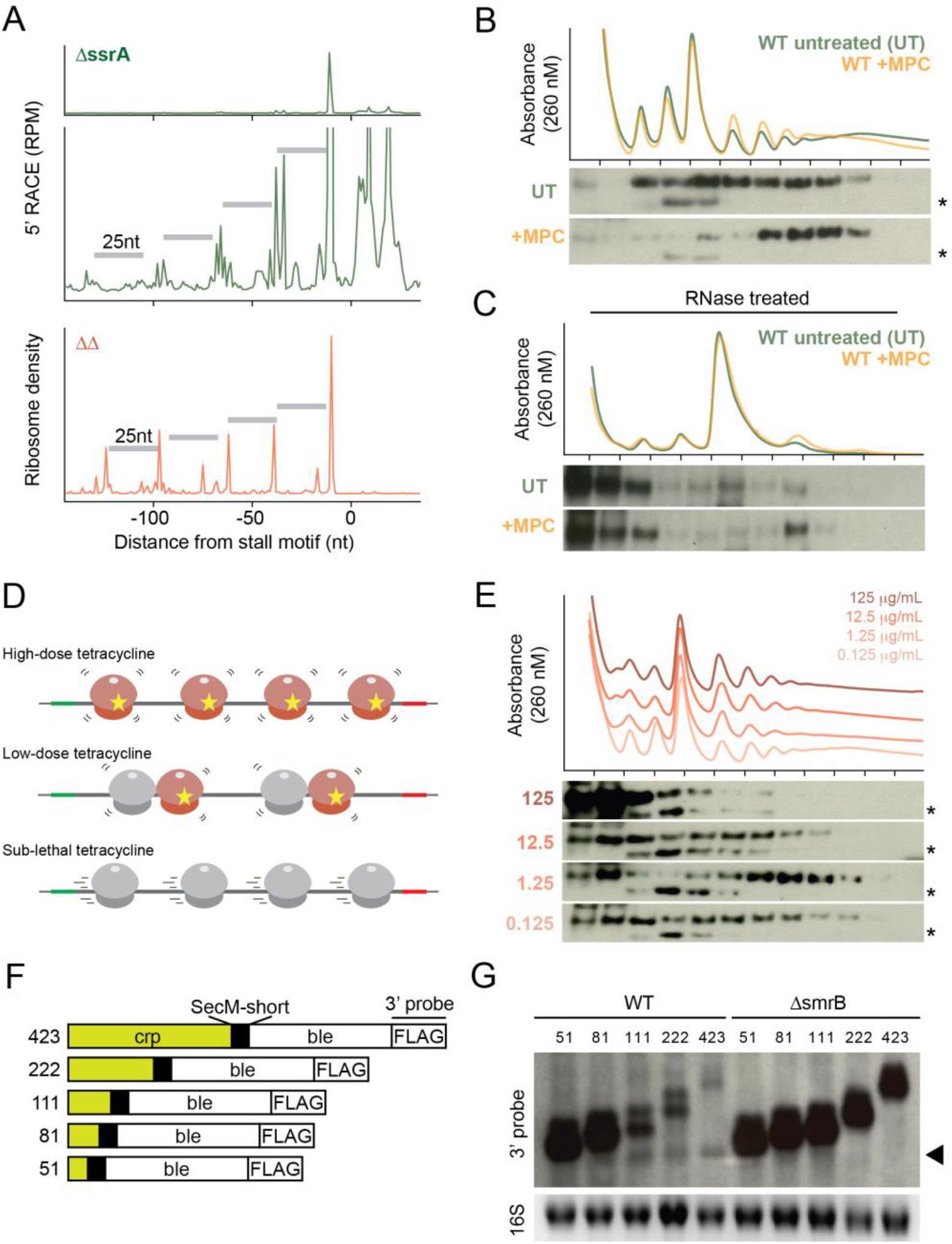
SmrB preferentially binds and cleaves upstream of collided ribosomes. (**A**) Top: 5’-RACE data reveal the SmrB cleavage sites as in Fig 4C, zoomed in to show smaller peaks upstream. Bottom: ribosome profiling data showing the 5’-boundary of ribosomes on the SecM reporter in the strain lacking tmRNA and SmrB. (**B**) The distribution of FLAG-SmrB with and without mupirocin (MPC) treatment (which induces pauses at Ile codons) was determined by fractionation over sucrose gradient and detection with an anti-FLAG antibody. A non-specific band is marked with *. (**C**) Lysates from cells with and without mupirocin treatment were treated with RNase A, fractionated over sucrose gradients, and the binding of FLAG-SmrB to nuclease resistant disomes was detected with an anti-FLAG antibody. (**D**) and (**E**) Low doses of tetracycline induce collisions whereas high doses stall ribosomes without inducing collisions. Following treatment with four different tetracycline concentrations, the distribution of FLAG-SmrB was determined by fractionation over sucrose gradient and detection with an anti-FLAG antibody. (**F**) and (**G**) In a new series of reporters, the Crp gene and bleomycin resistant gene are fused with the short SecM motif between them. The Crp gene is trimmed to reduce the number of ribosomes that can be loaded between the start codon and the stall site (the number shown). Reporter mRNA was detected on northern blots using the 3’-probe. An arrow indicates the downstream fragments. Ethidium bromide staining of 16S rRNA serves as a loading control.

Our work so far has focused on strong stalling motifs in reporter genes. To ask what effects collisions have on SmrB binding to ribosomes globally, we induced collisions throughout the transcriptome using the antibiotic mupirocin (MPC) which inhibits isoleucine tRNA synthetase and globally slows down the rate of decoding of Ile codons (*28, 31*). In untreated cells, FLAG-SmrB is distributed broadly across the sucrose gradient, extending from the subunit fractions to the polysome fractions (Fig 4B), arguing that most SmrB is ribosome-bound. After inducing collisions for 5 min with 50 μg/mL MPC, we observed that FLAG-SmrB moves deeper into the polysome fraction, consistent with preferential binding to collided ribosomes on messages that are heavily translated (Fig 4B).

Further, we treated cell lysates with RNase A to generate nuclease-resistant disomes, a hallmark of ribosome collisions (*17*). As expected, RNase A treatment collapses polysomes to yield a strong monosome peak and a small nuclease-resistant disome peak (Fig 4C). The disome peak is modestly but reproducibly higher in the MPC treated samples, consistent with the expectation that there are more ribosome collisions in MPC-treated cells. Although much of the SmrB dissociates from ribosomes under these conditions, moving into the top fractions, we see a strong SmrB band in the nuclease-resistant disome peak in the MPC-treated sample; quantitation of the amount of SmrB bound to various fractions shows strong enrichment for SmrB binding on colliding ribosomes relative to monosomes.

Another strategy to differentiate the effects of stalling and ribosome collisions is to treat cells with antibiotics that target the ribosome and then compare the effects of high doses, which stall all ribosomes quickly, versus lower doses that only stall some ribosomes, allowing others to translate until collisions occur (Fig 4D) (*18*). In untreated samples SmrB is broadly distributed in sucrose gradients while SmrB is enriched in polysomes deeper in the gradient when 1.25 μg/mL tetracycline is used (Fig 4E). Importantly, the enrichment of SmrB in the polysomes is lost in cells treated with 10- or 100-fold higher concentrations of tetracycline.

### Ribosome collisions promote SmrB cleavage

As previously performed in yeast (*18*), we generated a series of reporters to test whether ribosome collisions are required for mRNA cleavage. In these reporters, different lengths of the *crp* gene were fused upstream of the short SecM stalling motif, while the downstream *ble* sequence remains constant (Fig 4F). The reporters are numbered by the distance from the start codon to the stall site (in nt). We anticipated that the closer the short SecM motif is to the 5’-end of the ORF, the less room there is for ribosomes to load onto the mRNA and collide at the stalling motif. Using the 3’-probe against the reporter mRNA to follow the activity of SmrB, we observe the downstream mRNA fragment characteristic of SmrB cleavage and a strong reduction in full-length mRNA in the 111, 222, and 423 reporters (Fig 4G). In contrast, the downstream mRNA fragment is reduced in the 51 and 81 reporters and we see much higher levels of full-length mRNA. In the 51 and 81 reporters, only one or two ribosomes can be loaded upstream of the SecM stalled ribosome, respectively, suggesting that multiple collisions may be required to recruit SmrB. In the ΔsmrB strain, the amount of full-length mRNA is strongly increased in all five reporters and no downstream fragment is observed. These findings show that SmrB activity is triggered by collisions.

### The structure of collided ribosomes from B. subtilis and E. coli

Ribosomal collisions in yeast and mammalian cells create a distinct architecture of disomes with new composite interaction surfaces that are recognized by collision sensors (*16, 17*). To ask whether bacterial ribosomes display a similar behavior, we generated collided ribosomes in cell-free translation systems, translating mRNAs encoding the arrest peptides MifM (*32*) in *B. subtilis* extracts and VemP (*33*) in the commercially available *E. coli* PURE system. Following separation by sucrose density gradient centrifugation, disome and trisome peaks were collected and subjected to structural analysis by cryo-EM.

3D reconstruction of the *E. coli* disomes (details in Fig S5) revealed a defined arrangement of two ribosomes (Fig 5A). The leading (stalled) ribosome closely resembles the previously described VemP-stalled 70S in a non-rotated state, carrying a peptidyl-tRNA in the P site, a highly structured nascent peptide chain in the ribosomal tunnel, and an aminoacyl-tRNA in the A site (*34*). The collided ribosome is found in a mixture of rotated and non-rotated states carrying two tRNAs in the canonical or hybrid state conformation. The stalled ribosome engages the collided ribosome in an intricate interaction mainly involving protein-protein and protein-rRNA interactions between the two juxtaposed small 30S subunits. These interactions involve ribosomal proteins uS10, uS2, 16S rRNA helix h16, uS4, and 16S rRNA helices h5 and h17 in the collided ribosome interacting with uS9, uS2, 25S rRNA helix H78, uS11 and bS6, and uL9 of the stalled ribosome, respectively (Fig 5B,C). In addition, the L1 stalk in the large subunit of the stalled ribosome was observed in its “out” conformation forming a new bridge between its rRNA helix H78 and 16S helix h16 of the collided ribosome (Fig 5C).

**Figure 5.**
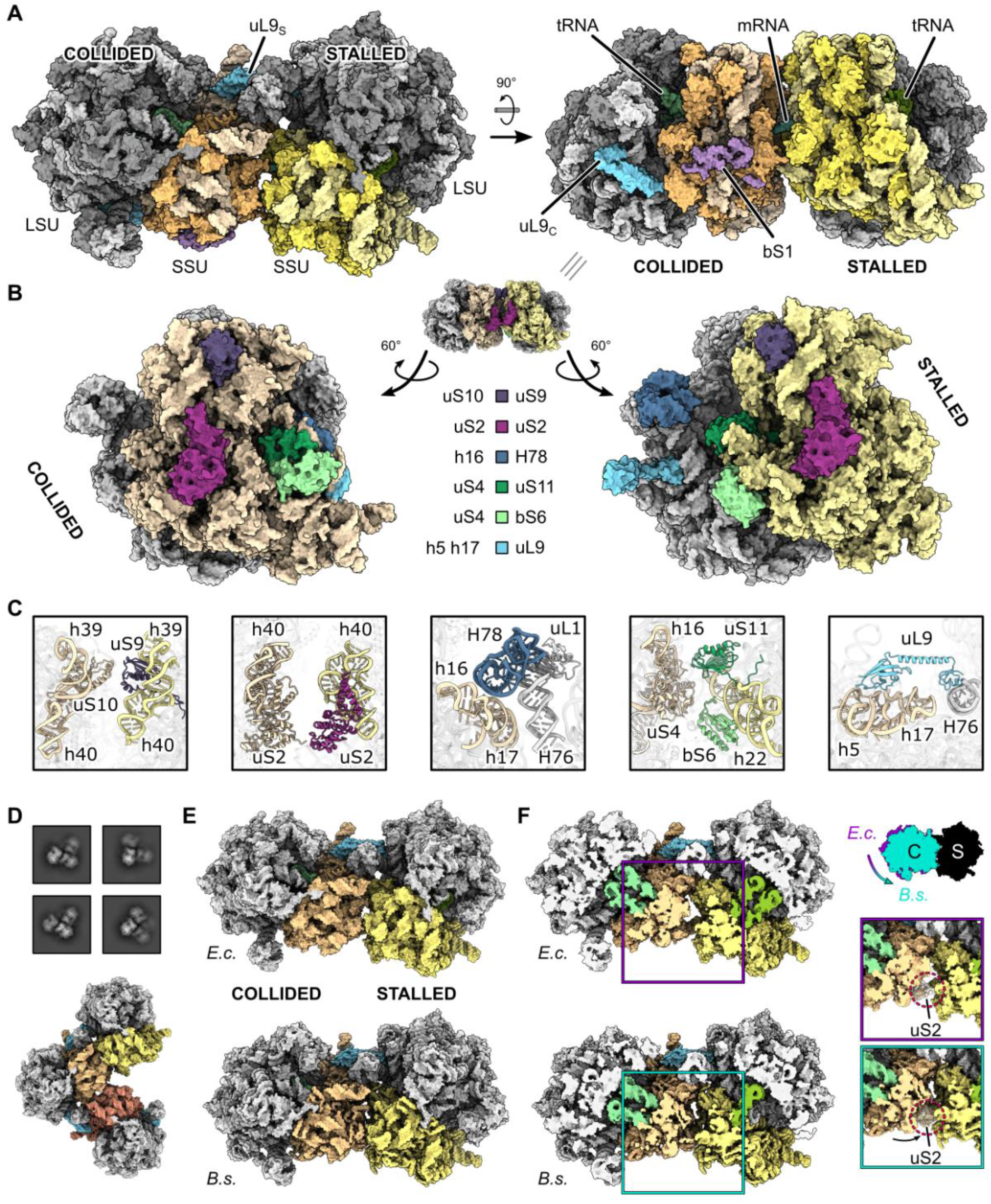
Cryo-EM structure of the *E. coli* disome. (**A**) Surface representation of the structural model of the *E. coli* disome. The uL9 proteins from stalled (uL9_S_) and collided (uL9_C_) adopt different conformations. (**B**) Interactions between stalled and collided ribosome at the disome interface. The disome interface is opened up by rotation of the stalled and collided ribosomes and interaction partners are shown in matching colors. (**C**) Cartoon representation of the individual interactions as they occur at the interface. (**D**) 2D class averages and cryo-EM structure model of an *E. coli* trisome. (**E** and **F**) Comparison of the *E. coli* (*E.c.*) and *B. subtilis* (*B.s.*) disomes displaying full and cut views. Note the smaller space between stalled and collided ribosomes in the *B.s.* disome interface as illustrated by comparing the positions of uS2 proteins in the zoomed view in (F).

Another substantial difference between the conformation of the stalled and collided ribosome is the dramatic rearrangement of the uL9 protein of the large ribosomal subunit. In individual 70S ribosomes, from its binding site on the 50S subunit below the L1 stalk, L9 contacts uS6 and uL2 in the 30S subunit (not shown). While this conformation was also observed in the collided ribosome, the L9 protein of the stalled ribosome flipped out of this position to engage in a novel mode of interaction via its C-terminal domain with the 30S subunit of the collided ribosome. This interaction involves rRNA helices h5 and h17 of the collided ribosome, thereby effectively forming another bridge between the stalled 50S subunit and the collided 30S subunit (Fig 5C). Although a similar bridging interaction of L9 between individual 70S ribosomes was observed previously in crystallized 70S ribosomes from *E. coli* and other bacteria (*35, 36*), the L9 binding site on the neighboring 30S subunit does not overlap with the site observed here in collided disomes and may be an artifact caused by crystallization conditions.

Notably, the largest protein of the 30S subunit, S1, is missing in the stalled ribosome but present in the collided one (Fig 5A). The arrangement of the interface between the solvent sides of the two 30S subunits results in very limited accessible space and would lead to a steric clash of the S1 protein with the collided ribosome. We conclude that formation of the observed disome architecture requires dissociation of S1 from the stalled ribosome. S1 dissociation may serve as a checkpoint in order to discriminate between short-lived ribosome collisions in productive polysomes and longer lasting stalling events.

When analyzing *E. coli* trisomes, we found that the structural features of the interface between the second and third ribosomes are essentially identical to the interface observed between the stalled and first collided ribosomes (e.g. the L1 and L9 bridges) (Fig 5D). This suggests that during long-lived stalling events, additional collisions can accumulate and yield a multitude of composite ribosome-ribosome interfaces. This may explain our observation of gradually increasing efficiency of ribosome rescue with longer reporter mRNAs allowing for more ribosomes to collide (Fig 4F,G).

We wondered whether the observed disome formation is unique to *E. coli* and compared our structures to those of the MifM-stalled disomes from the Gram-positive bacterium *B. subtilis*. We found that ribosome collisions in *B. subtilis* result in disomes adopting an essentially identical conformation as observed in *E. coli*: the overall orientation and interactions of the ribosomes are highly similar, the bridge by the L1 stalk to rRNA helix h16 is formed, and L9 of the stalled ribosome reaches over to the 30S subunit of the collided one (Fig 5E). One notable difference is that the 30S subunits are positioned closer in the *B. subtilis* disome when compared to *E. coli* (Fig 5F). This high degree of overall similarity indicates that the observed mode of collided disome formation is likely to be conserved in bacteria, with subtle differences at the disome interface. Notably, the overall architecture of these collided disomes is very different from hibernating, so-called 100S disomes formed under stress conditions (Fig S6) (*37, 38*). We conclude that, similar to eukaryotes, the observed disome (and trisome) architecture of collided ribosomes is a conserved feature in bacteria that can provide a unique interface used by rescue factors such as SmrB to recognize stalled ribosomes.

### In vitro cleavage of mRNA and the SmrB-disome structure

Next, we reconstituted SmrB recruitment to collided disomes and endonucleolytic cleavage of mRNA *in vitro*. We incubated purified SmrB or the nuclease deficient ALA mutant with purified VemP-stalled disomes and analyzed the reaction products by sucrose density gradient centrifugation (Fig 6A). In a control reaction without SmrB we observed disomes, as expected, as well as some 70S ribosomes likely arising from ribosome dissociation from the mRNA and/or background nucleolytic activity. In contrast, after incubation with wild-type but not mutant SmrB, we observed an almost complete loss of the disome signal and a corresponding increase in the 70S signal, consistent with cleavage between the collided ribosomes by the endonuclease activity of SmrB.

**Figure 6.**
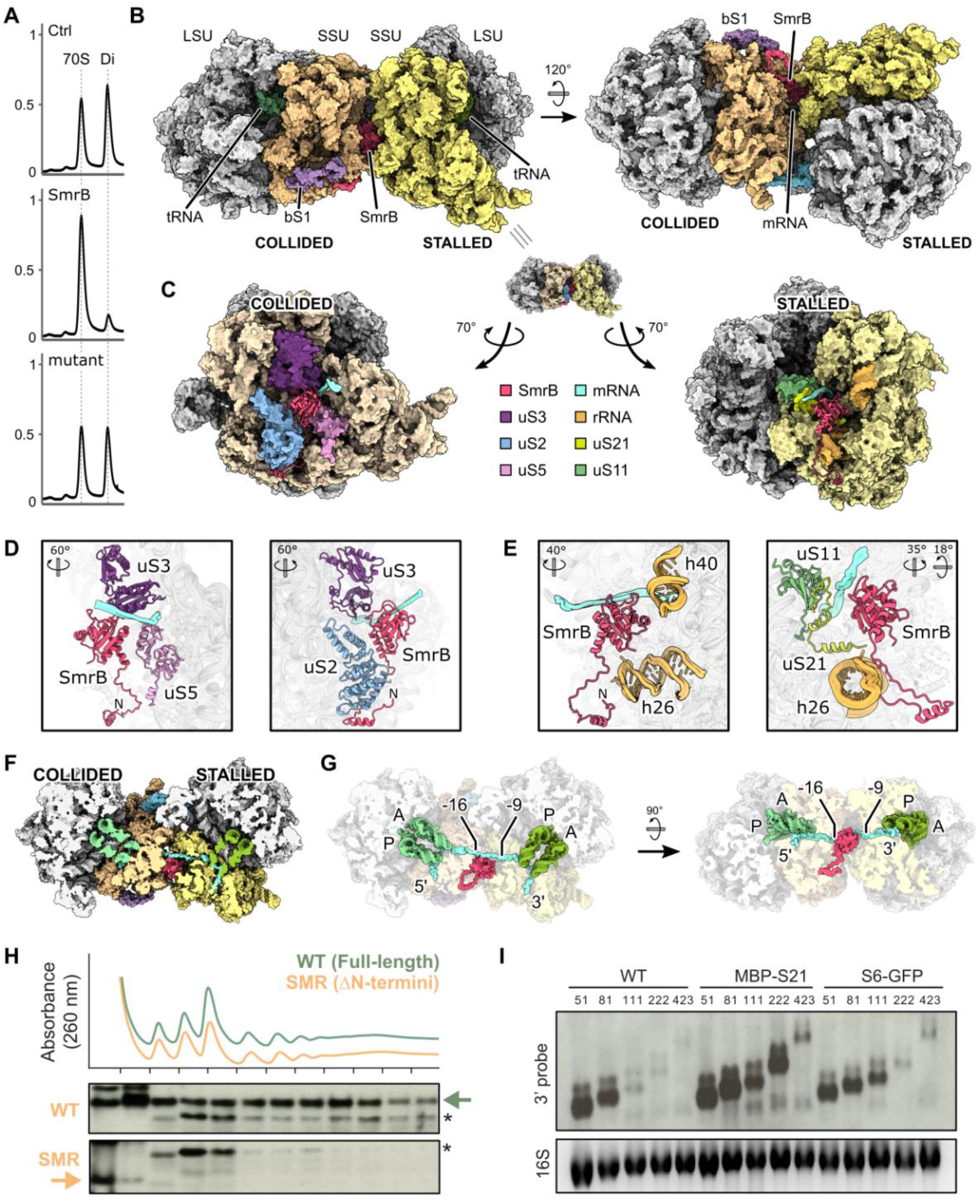
Cryo-EM structure of the SmrB-bound *E. coli* disome. (**A**) Disome nuclease assay. Sucrose density gradient profiles of VemP-stalled disomes alone (Ctrl), incubated with SmrB (SmrB) and incubated with nuclease deficient SmrB (mutant), respectively. The Y-axes show the absorption at 260 nm. (**B**) Surface representation structural model of SmrB bound to the *E. coli* disome. (**C**) Interactions between SmrB and the stalled and collided ribosomes at the disome interface. The disome interface is opened up by rotation of the stalled and collided ribosomes and SmrB is shown in cartoon representation. (**D** and **E**) Cartoon representation of the individual interactions of SmrB with the collided ribosome in (D) and the stalled ribosome in (E). The orientations of the individual views with respect to (C) are indicated. (**F**) Cut view of the SmrB bound disome showing the mRNA path. (**G**) Interaction of SmrB with the mRNA. The disome orientation in the left part of (G) corresponds to the one shown in (F). Approximately 8 nucleotides of the mRNA are exposed at the disome interface, in reach for SmrB cleavage. The nucleotides at the ribosome boundaries are indicated, counting the first nucleotide in the A-site of the stalled ribosome as 0. (**H**) The distribution of FLAG-tagged full-length SmrB and a construct with only the SMR domain (residues 88-183) was determined by fractionation over sucrose gradient and detection with an anti-FLAG antibody. A non-specific band is marked with *. (**I**) Northern blots using the 3’-probe against the CRP reporters with the short SecM stalling motif in wild-type cells, a strain where MBP is fused to the N-terminus of S21, and a strain where GFP is fused to the C-terminus of S6.

We reconstituted the nuclease-deficient SmrB mutant with our *E. coli* VemP-stalled disomes and subjected the complexes to structural analysis by cryo-EM (details in Fig S7). Compared to disomes alone, this reconstruction revealed an extra density between the 30S ribosomal subunits in the immediate vicinity of the mRNA stretching from the mRNA exit site of the stalled ribosome to the mRNA entry of the collided ribosome (Fig 6B). Although the overall resolution of the SmrB-bound disome is 3.3 Å, we observed limited local resolution in this region of the map. Therefore, we could only partially build the molecular structure of SmrB and relied on the AF2-driven prediction for rigid body docking of SmrB (*39*) (Fig S8).

The binding site for SmrB involves both the stalled and the collided ribosome: on the collided ribosome the SMR domain interacts with uS3, uS2 and uS5, whereas on the stalled ribosome it binds rRNA helices h40 and h26 as well as ribosomal proteins uS11 and uS21 (Fig 6C–E). Notably, in several bacteria the genes encoding uS21 and SMR-domain proteins are tightly linked in conserved operons, suggesting that this interaction might be a conserved aspect of binding of SMR domains to collided ribosomes. The N-terminal extension region of SmrB forms a hook-like structure that wraps around uS2 of the collided ribosome (Fig 6D). Interestingly, the N-terminal alpha-helix contains the --xxxa motif conserved in other SMR-domain proteins in proteobacteria (Fig 2C and Fig S3). We observed this hook-like N-terminus of SmrB on the stalled ribosome as well, although in that position it cannot be connected to the SMR domain between the subunits and thus represents a second copy of SmrB in the complex. We speculate that this N-terminal helix may promote the initial recruitment of SmrB to elongating 70S ribosomes and that when collisions occur, the subsequent binding of the SMR domain at the composite binding site between collided ribosomes further stabilizes SmrB binding. In agreement with this idea, a truncated SmrB mutant consisting of only the SMR domain (residues 88-183) does not bind ribosomes (Fig 6H). The observed binding mode of SmrB therefore explains how SmrB is specifically recruited to collided disomes (or trisomes).

The active site of SmrB interacts with the bridging mRNA, poised for cleavage in between the individual ribosomes (Fig 6F,G). From the structure, it is difficult to determine the exact mRNA residues to be cleaved, however, and the nuclease-deficient mutant of SmrB may engage in a somewhat different interaction with its mRNA substrate. Nevertheless, the observed positioning indicates that SmrB can execute endonucleolytic cleavage of the mRNA between position −9 and −16, counting from the first nucleotide in the A site of the stalled ribosome, in agreement with our biochemical data. We speculate that activation of SmrB specifically on collided disomes is a result of precise positioning with respect to its substrate.

### Disruption of the SmrB-binding pocket on collided ribosomes

We asked if disruption of the disome interface or the SmrB binding pocket formed between collided ribosomes would interfere with mRNA cleavage by SmrB. The L9 protein from the stalled ribosome makes contacts with the 30S subunit in the collided ribosome (Fig 5C) and strains lacking L9 are known to have high levels of frameshifting (*40, 41*). We found, however, that SmrB still cleaves the CRP reporters in a collision dependent manner in an L9 knockout strain; the reporter mRNA processing is indistinguishable from that seen in wild-type strains (Fig S9). Moreover, fusion of mCherry to the C-terminus of L9 (on the domain that contacts 16S rRNA in the collided ribosome) also has no effect on SmrB activity (Fig S9). These results suggest that this contact between the disomes does not play an essential role in stabilizing the disome interaction. In contrast, we observe that fusion of MBP to the N-terminus of S21 dramatically stabilizes full-length mRNA in the 111 and 222 CRP reporters compared to the wild-type strain (Fig 6I). To a lesser extent, fusion of GFP to the C-terminus of S6 also stabilizes the 111 CRP reporter compared to the wild-type strain (Fig 6I). These results are consistent with a reduction in SmrB activity and stabilization of the reporter mRNA due to a disruption of the SmrB binding pocket formed between the collided ribosomes, validating the structural findings reported here.

## Discussion

Our findings indicate that ribosome collisions are the critical trigger for ribosome rescue in *E. coli*: collisions recruit the endonuclease SmrB, triggering mRNA cleavage and rescue of upstream ribosomes by tmRNA. Under normal conditions, SmrB is distributed broadly across sucrose gradients, suggesting that it binds ribosomes generally, scanning for problems. When collisions are induced throughout the transcriptome, SmrB moves deep into the polysome fraction and binds preferentially to nuclease-resistant disomes that are a hallmark of collisions. Importantly, we find that SmrB recruitment is triggered by low doses of tetracycline that promote collisions but not by high doses that promote stalling without collisions. Furthermore, SmrB is unable to cleave mRNA at stalling motifs positioned too close to the 5’-end of an ORF for sufficient collisions to occur. We conclude that ribosome collisions serve as a signal to recruit ribosome rescue factors in bacteria as well as in eukaryotes.

The importance of collisions is fully validated by our cryo-EM structures of collided ribosomes from both Gram negative *E. coli* cells and Gram positive *B. subtilis* cells that reveal a specific architecture of closely interacting individual 70S ribosomes. Their architecture is similar to, but distinct from, the collided disome structures previously characterized in eukaryotic yeast and mammalian cells (*16, 17*). In both cases, the interaction between stalled and collided ribosomes primarily involves the two small ribosomal subunits, but also employs contacts between the large subunit of the stalled ribosome and the small subunit of the collided one. However, these contacts are not conserved between the kingdoms: in eukaryotes they are established by 25S rRNA helix H31L and ribosomal protein eL27 of the large subunit, whereas in bacteria by the L1 stalk and the ribosomal protein uL9. The collided disome architecture is likely to be conserved in bacteria and, importantly, differs completely from the structure of hibernating disomes in *E. coli* and *B. subtilis*. This makes the observed collided disomes a valid proxy for sensing ribosome stalling in bacteria.

A role for collisions in ribosome rescue is consistent with previous studies in bacteria. Structural studies of polysomes and the crystal packing interactions in x-ray structures show that bacterial ribosomes pack closely together through interactions between their small subunits (*35, 42*). Moreover, in ribosome profiling studies in *E. coli*, Subramaniam et al. observed reduced ribosome density downstream of pausing sites due to the removal of stalled ribosomes by tmRNA (*43*); these researchers later argued that reductions in protein output were most consistent with a mathematical model in which collisions are the trigger that recruits rescue factors to remove stalled ribosomes (*44*). Finally, two recent studies suggest that ribosome collisions influence the level of frameshifting at pause sites in *E. coli* (*45, 46*).

It is striking that collided ribosomes are recognized in *E. coli* by an SMR-domain protein (SmrB) given that the same domain plays a similar role in yeast (Cue2) (*20*). Prior to this study, only Rqc2, the factor that promotes CAT-tailing in yeast and has homologs in some bacteria (*47, 48*), has been shown to mediate ribosome rescue pathways in both bacteria and eukaryotes. Our analysis identified SMR-domain proteins in all but a few bacterial phyla (Fig 2A), suggesting that a role for SMR proteins in ribosome rescue may be widespread in bacteria just as it is in eukaryotes.

In yeast (*20*), worms (*21*), and now *E. coli*, ribosome collisions lead to mRNA cleavage by SMR-domain proteins upstream of the ribosome stalling site, targeting the problematic mRNA for decay. Cleavage leads to the rescue of upstream ribosomes which, no longer impeded by a downstream stalled ribosome, translate to the 3’-end of the upstream fragment and are released by rescue factors, tmRNA-SmpB in bacteria and Dom34-Hbs1 in yeast. Despite these general similarities, there are also a few apparent differences between the activity of SmrB and the Cue2 protein in yeast. While our RACE and MS data place the SmrB cleavage site at the 5’ boundary of the stalled ribosome (Fig 3), Cue2 cleavage has been mapped to the A site of the collided ribosome in a wild-type yeast strain and to the 5’ boundary of the stalled ribosome in strains lacking rescue factors Hel2 or Slh1 (*16, 20, 49*). A second difference is that loss of Cue2 alone has little or no effect on the stability of reporter mRNAs in yeast because the processive 5’-3’ exonuclease Xrn1 is primarily responsible for their decay (*20*). In contrast, loss of SmrB leads to a dramatic increase in full-length mRNA in our reporters in *E. coli* (Fig 2F).

We observe two pathways by which stalled ribosome complexes are resolved (Fig 3I). The main pathway is cleavage at the 5’-boundary of the stalled ribosome by SmrB. In the second pathway, 3’-to-5’ exonucleases degrade mRNA until they encounter the stalled ribosome, allowing tmRNA access for rescue to occur (*30, 50*). *E. coli* possesses three major 3’-to-5’ exonucleases involved in mRNA turnover: RNase II, RNase R, and polynucleotide phosphorylase. Previous studies showed that strains with single knockouts of any of these exonucleases still exhibited trimming of mRNA back to the 3’-boundary of ribosomes stalled at the SecM motif (*50*); the triple deletion strain is not viable.

The structure of the disome-bound SmrB illustrates how this rescue factor can generally screen translating ribosomes and, in case of problematic events such as mRNA damage by oxidative or alkylating agents (*3*), is able to specifically recognize the new composite interface formed between the individual collided 70S ribosomes. Through its N-terminus, SmrB associates generally with elongating ribosomes for screening, whereas the composite binding site between the adjacent collided 30S subunits is required to position the SMR domain of SmrB proximal to the bridging mRNA for endonucleolytic cleavage. In contrast, eukaryotic SMR-domain proteins frequently contain ubiquitin-binding domains (e.g. CUE, UBA, UIM, and UBL) to bind ribosome proteins ubiquitinylated by E3 ligases such as Hel2. This diversity suggests that recruitment to collided ribosomes can occur through multiple alternate mechanisms.

In this report, we reveal a novel ribosome rescue factor in bacteria, SmrB, that recognizes ribosome collisions and specifically targets stalled ribosomes for mRNA decay and ribosome rescue. Bacteria and eukaryotes both rely on collisions to sense ribosome stalling and SMR-domain proteins to cleave the mRNA so that upstream stalled ribosomes can then be rescued by factors known to act on truncated mRNAs. These common features substantiate the universal significance of ribosome collisions and endonucleolytic cleavage in ribosome rescue. We anticipate the bacterial system to be as complex as the eukaryotic counterpart that enables recognition of stalled ribosomes, triggering RNA and proteolytic processing, and rescuing ribosomes. Our present work provides the initial scaffold for additional studies to fully appreciate this phenomenon.

## Supporting information

List of strains, plasmids, and oligonucleotides

## Acknowledgements

The authors thank Suparna Sanyal for sharing *E. coli* strains QC101 and QC901, Haiping Hao at the JHMI Transcriptomics and Deep Sequencing Core for assistance with high-throughput sequencing, Bob Cole and Tatiana Boronina at JHMI in the Mass Spectrometry and Proteomics Facility, Joanna Musial for assistance during protein purifications, C. Ungewickell and S. Rieder for technical assistance, and L. Kater and K. Best for support with the pre-processing pipeline of cryo-EM data.

## Author contributions

K.S. performed the genetic screen, the analyses of the nanoLuc-ble reporters, and sucrose gradients. A.C. analyzed the CRP reporters and prepared samples for the MS experiments. M.B. and L.A. performed the phylogenetic analyses. H.K. performed in vitro translation and in vitro nuclease assays and prepared samples for cryo-EM analysis. O.B. and R.Bu. collected cryo-EM data. H.K. processed the cryo-EM data and R.Bu. prepared molecular models. H.K., R.Bu. and R.Be. analyzed and interpreted the structures and R.Bu. prepared structural figures. A.B., R.G., and R.Be. supervised the project.

## Data and materials availability

The ribosome profiling data are available at the GEO with accession number GSE179691. Cryo-EM volumes and molecular models have been deposited at the Electron Microscopy Data Bank and Protein Data Bank with accession codes EMD-XXXX (*E. coli* maps) and PDB-YYYY (stalled 70S) PDB-ZZZZ (collided 70S-SmrB), EMD-YYYY (*B. subtilis* maps).

