## Supplemental Methods and Figures for "Ribosome collisions in bacteria promote ribosome rescue by triggering mRNA cleavage by SmrB"

### SUPPLEMENTARY INFORMATION

#### MATERIALS & METHODS

##### Bacterial strains and plasmids

A list of strains and plasmids and the details of their construction are given in Table S1.

The L9 knockout strain (BW25113 rplI::kan) was obtained from the *E. coli* Genetic Stock Center. Strains QC101 and QC901 containing the L9-mCherry and S6-GFP, respectively, were gifts from Suparna Sanyal (*51*). Additional knockout strains of MG1655 were constructed using one-step genomic replacement with a PCR fragment with λ Red recombinase as described (*52*). Gene deletions and the endogenous epitope-tagged SmrB mutants were verified by PCR and sequencing.

The nanoLuc-ble reporter construct pKS-nonstall was expressed from plasmids containing an AmpR marker and a p15A origin of replication. DNA encoding various stalling motifs was inserted between the genes for nanoLuc and ble using Gibson assembly.

To construct the Crp collision reporters, the first 39, 69, 99, 109, and 411 bases of the *crp* coding region were amplified from MG1655 genomic DNA. The PCR also added the sequence encoding the short SecM ribosome stalling motif (GIRAGP) in-frame immediately downstream of the *crp* fragment. The PCR products were inserted into EcoRI and BglII digested pKS-nonstall to produce the pCRP51, pCRP81, pCRP111, pCRP121, and pCRP423 reporter plasmids using Gibson assembly. The names of the plasmids and numbers in the text represent the distance from the AUG to the stall site (the second Gly codon in GIRAGP) in each reporter.

To construct the plasmid pAC01 encoding tmRNA-DD, we switched the origin of replication of the pKW23 plasmid (*4*) from p15A to pBR322 for compatibility. This was accomplished by Gibson assembly of two PCR products: 1) everything in pKW23 except the origin and 2) the pBR322 origin from pBAD-GFPuv (*53*).

For overexpression and purification of SmrB, we amplified the *smrB* gene from genomic DNA from MG1655, adding a Twin-Strep tag, a TEV cleavage site, and a FLAG tag to the N-terminus of SmrB using nested PCR primers. This amplicon was inserted into pET24b cleaved with NdeI and BamHI using Gibson assembly.

##### Genetic screening

A library with random transposon insertions throughout the genome was constructed using the EZ-Tn5 <KAN-2>Tnp Transposome Kit (Lucigen). The EZ-Tn5 transposome was electroporated into the parental *E. coli* strain MG1655 and the cells were plated on LB + kanamycin (50 mg/L). After one overnight incubation, approximately 5 million colonies were collected and stored at –80 °C in LB + 20% glycerol. Subsequently, either the SecM or Ile8 reporter plasmid was electroporated into the random insertion library and transformants were plated on LB + ampicillin (50 mg/L) + phleomycin (50 mg/L). After one overnight incubation, phleomycin resistant colonies were collected and stored in at –80 °C in LB + 20% glycerol.

To quantify the number of transposon insertions in each gene throughout the genome, Tn-seq was performed on the random insertion library as well as the libraries of phleomycin-resistant colonies from both the SecM and Ile8 screens. Sequencing libraries were prepared from genomic DNA from each library using the NEBNext Ultra II FS DNA Library Prep Kit for Illumina (NEB) following the manufacturer's protocol. During PCR amplification of the adapter-ligated DNA, we use two PCR steps with primers that specifically bind to the mosaic end of EZ-Tn5 <KAN-2> Transposon to prepare libraries enriched in transposon insertion sites. Between the PCR steps, we included an additional enrichment step based on biotin-streptavidin purification. The two-PCR steps were performed using NEBNext Ultra II Q5 Master Mix (NEB) as follows: The 1st PCR enrichment was performed with primers KS_Tn5_1stPCR_F_biotin and KS_Tn5_1stPCR_R (Table_S1) with the following program: 30 s at 98 °C; 15 cycles of 10 s at 98 °C, 20 s at 59 °C, 60 s at 65 °C; and 5 min at 65 °C. The PCR products were purified first with DNA Clean & Concentrator-5 columns (Zymo Research) and then Dynabeads MyOne Streptavidin C1 beads (Thermo Fisher). The streptavidin beads were washed four times with binding and washing buffer (5 mM Tris-HCl pH 7.5, 0.5 mM EDTA pH 8.0, 1M NaCl); the PCR products were added and incubated for 30 min at 25 °C; and the beads were washed twice with the binding and washing buffer and then twice by 0.1x TE buffer (1 mM Tris pH 8.0, 0.1 mM EDTA pH 8.0). The beads bound to the PCR products were used directly as the template for the 2nd PCR enrichment using KS_Tn_library_F and one of our custom Index primers for Tn-seq (Table S1) with the following program: 30 s at 98 °C; 7 ~ 9 cycles of 10 s at 98 °C, 20 s at 59 °C, 60 s at 65 °C; and 5 min at 65 °C. The products from the 2$^{nd}$ PCR were gel purified on a non-denaturing 5% TBE gel, analyzed on a BioAnalyzer high sensitivity DNA kit (Agilent), and sequenced on the NextSeq 500 instrument (Illumina).

The Tn-seq data were analyzed with custom scripts written in Python 2.7. The adaptor sequence AGATCGGAAGAGCACACGTC was removed from the 3'-ends of reads with Skewer (*54*). Reads of interest contain the Tn5 transposase sequence at the 5'-end and genomic DNA sequence at the 3'-end. Reads lacking the Tn5 transposase sequence GGTTGAGATGTGTATAAGAGACAG at their 5'-ends were discarded using cutadapt. After trimming this sequence, the remaining reads were aligned to *E. coli* MG1655 genome build NC_000913.2 using bowtie version 1.1.2 (*55*). The site of the transposon insertion was assigned using the 5'-end of the aligned reads. The number of transposon insertion sites for each gene was counted and normalized as reads per kilobase per million mapped reads (RPKM), normalizing for both the sequencing depth for each library and the length of each gene.

**Sequence Analyses**

PSI-BLAST (*56*) and JACKHMMER programs (*57*) were used to carry out iterative sequence profile searches to collect Smr domain-containing sequences. Proteins were clustered using BLASTCLUST (https://ftp.ncbi.nih.gov/blast/documents/blastclust.html) to identify shared domain architectural themes. Additional domains fused to the Smr domain were annotated using a database of domain sequence profiles including pfam A models (*58*). For contextual analysis of prokaryotic gene neighborhoods, the GenBank genome files corresponding to unique GenBank genome assemblies (GCA ids) were used as starting material. Specific neighborhoods were extracted using a Perl script that reports upstream and downstream genes of the anchor Smr domain-containing gene. Proteins encoded by these genes were then clustered using BLASTCLUST to identify conserved gene neighborhoods based on conservation between different taxa. Additional filters outputted valid neighborhoods for further analysis: (1) nucleotide distance constraint (generally 50 nucleotides), (2) conservation of gene directionality within the neighborhood, and (3) presence in more than one phylum. Multiple sequence alignments were built using the Kalign program (*59*), and manually improved based on the alignments outputted by sequence homology searches. Secondary structure prediction was done using the JPred program (*60*). Phylogenetic relationships were determined using an approximate maximum likelihood (ML) method as implemented in the FastTree program (*61*): corresponding local support values were also estimated as implemented. To increase accuracy of topology, the rounds of minimum-evolution subtree-prune-regraft (SPR) moves were increased to 4 (-spr 4) and we utilized options -mlacc and -slownni to survey more exhaustively the ML nearest neighbor interchanges (NNIs).

**Western blots**

Cells were grown in LB + ampicillin (50 mg/L) to $OD_{600}$ = 0.5, harvested by centrifugation, resuspended in 12.5 mM Tris pH 6.8 with 4% SDS, and lysed by heating to 90 °C for 10 min. 5x loading dye (250 mM Tris pH 6.8, 20%

glycerol, 30% β-mercaptoethanol, 10% SDS, saturated bromophenol blue) was added and the lysate was denatured at 90 °C for 10 min. Protein was separated on a 4–12% Criterion XT Bis-Tris protein gel (Bio-Rad) using XT MES buffer and transferred to PVDF membrane using the Trans-Blot Turbo Transfer system (Bio-Rad). Membranes were blocked in 5% milk for 1 h at room temperature, washed, and then probed with antibodies diluted in TBS-tween. Antibodies dilutions were: anti-FLAG-HRP 1:10000 (Sigma); anti-Strep•Tag II-HRP 1:5000 (Millipore Sigma); anti-rpoB 1:1000 (BioLegend); anti-rpoC 1:1000 (BioLegend); and anti-mouse-HRP 1:2000 (Thermo Fisher). Chemiluminescent signals from HRP were detected using SuperSignal West Pico PLUS Chemiluminescent Substrate (Thermo Fisher) or SuperSignal West Femto Maximum Sensitivity Substrate (Thermo Fisher) and visualized on Amersham Hyperfilm ECL (GE).

**Northern blots**

Cells were grown in LB + ampicillin (50 mg/L) to $OD_{600}$ = 0.5, harvested by centrifugation, and resuspended in 100 mM NaCl, 10 mM Tris pH 8.0, 1 mM EDTA pH 8.0, and 1% SDS. RNA was extracted twice by phenol, pH 4.5 (once at 65°C and once at room temperature) followed by chloroform extraction. RNA in the aqueous layer was then precipitated by isopropanol and 0.3 M NaOAc (pH 5.5), washed with 80% ethanol, and resuspended in water. Purified RNA was separated on a 1.2% agarose-formaldehyde denaturing gel and transferred to a nylon membrane (Hybond-N+, Cytiva) in 10 x SSC buffer using a Model 785 Vacuum Blotter (Bio-Rad). RNA was crosslinked to the membrane with the Stratalinker UV crosslinker (Stratagene). Pre-hybridization and hybridization was performed in PerfectHyb Plus Hybridization Buffer (Milipore Sigma). RNA was probed with 50 nM 5'-digoxigenin labeled DNA oligos (IDT). Digoxigenin was detected with anti-Digoxigenin-AP antibodies diluted 1:1000 (Milipore Sigma). Chemiluminescent signals from alkaline phosphatase were detected with CDP-Star (Milipore Sigma) and visualized on Amersham Hyperfilm ECL (GE).

**5'-RACE**

RNA was extracted as described above. DNA contamination was depleted by treatment with RQ1 DNase (Promega). 5 μg of purified RNA was 5'-phosphorylated by incubating with T4 polynucleotide kinase (NEB) in 1 mM ATP at 37 °C for 30 min, after which PNK was denatured by heating to 75 °C for 10 min. The RNA adapter KS_5RACE_linker was ligated to the 5'-end of the RNA by incubating with T4 RNA ligase 1 (NEB) in 1 mM ATP and 15% PEG8000 at 25 °C for 3 h. Ligated samples were purified by 2.2x volume of RNAclean XP (Beckman). The 1st strand cDNA was synthesized using the KS_5RACE_RT primer and SuperScript III Reverse Transcriptase (Thermo Fisher) by incubating at 54 °C for 60 min after which RT was denatured by heating to 85°C for 5 min. Denatured reverse transcription products were used directly in the 1st PCR reaction. The 1st PCR was performed with Phusion High-Fidelity DNA Polymerase (NEB) and a set of primers of KS_5RACE_F1 and KS_5RACE_R1; with the program: 30 s at 98 °C; 25 cycles of 10 s at 98 °C, 10 s at 65 °C, 60 s at 72 °C; and 5 min at 72 °C. The 1st PCR products were purified using DNA Clean & Concentrator-5 columns (Zymo Research). The 2nd PCR was performed with Phusion High-Fidelity DNA Polymerase (NEB) and a set of primers of NI-NI-2 and one of our custom Index primers for 5'RACE (Table S1) with the program: 30 s at 98 °C; 25 cycles of 10 s at 98 °C, 10 s at 65 °C, 60 s at 72 °C; and 5 min at 72 °C. The 2nd PCR products were purified using DNA Clean & Concentrator-5 columns (Zymo Research), analyzed on a BioAnalyzer high sensitivity DNA kit (Agilent), and sequenced on the MiSeq Nano instrument (Illumina).

The 5'-RACE data were analyzed using custom scripts written in Python 2.7. The RNA adapter sequence TGCCCGAGTG was removed from the 5'-end of reads using cutadapt. Reads without the RNA adapter sequence were discarded. The reverse primer sequence GCGGTCGAGTTCTGGACCGA from the 2nd PCR was removed from the 3'-ends of reads by cutadapt. The processed reads were aligned to the SecM or EP* reporter plasmid sequences using bowtie version 1.1.2 (*55*). The 5' ends of mapped reads were counted and normalized as reads per million mapped reads (RPM), normalizing for the sequencing depth of each library.

**3'-RACE**

RNA was extracted as described above. DNA contamination was depleted by treatment with RQ1 DNase (Promega). 5 μg of purified RNA was 3'-dephosphorylated by incubating with T4 polynucleotide kinase (NEB) without ATP at 37 °C for 30 min, after which PNK was denatured by heating to 75 °C for 10 min. The 3' DNA adapter was 5' adenylated using the 5´ DNA Adenylation Kit (NEB) and purified with an Oligo Clean & Concentrator column (Zymo Research). This adaptor was ligated to 3'-end of dephosphorylated RNA by incubating with T4 RNA ligase 2 truncated (NEB) in 15% PEG8000 at 37 °C for 3 hours. Ligated samples were purified with RNAclean XP (Beckman). The 1st strand cDNA was synthesized with KS_3RACE_RT and SuperScript III Reverse Transcriptase (Thermo Fisher) by incubating at 54 °C for 60 min, after which RT was denatured by heating to 85°C for 5 min. Denatured reverse transcription products were used directly in the 1st PCR reaction. The 1st PCR was performed with Phusion High-Fidelity DNA Polymerase (NEB) and primers KS_3RACE_F1 and KS_3RACE_RT with the program: 30 s at 98 °C; 9 cycles of 10 s at 98 °C, 10 s at 65 °C, 60 s at 72 °C; and 5 min at 72 °C. The PCR products were purified using DNA Clean & Concentrator-5 columns (Zymo Research). The 2nd PCR was performed with Phusion High-Fidelity DNA Polymerase (NEB) and one of our custom Index primers for 3'RACE (Table S1) and KS_3RACE_R with the program: 30 s at 98 °C; 25 cycles of 10 s at 98 °C, 10 s at 65 °C, 60 s at 72 °C; and 5 min at 72 °C. The 2nd PCR products were purified by DNA Clean & Concentrator-5 columns (Zymo Research), analyzed on a BioAnalyzer high sensitivity DNA kit (Agilent), and sequenced on the MiSeq Nano (Illumina).

The 3'-RACE data were analyzed by custom scripts written in Python 2.7. The DNA adapter sequence TCCTTGGTGCCCGAGTGNNNNNN was removed from the 5'-end of reads using cutadapt. Reads without the adapter sequence were discarded. The forward primer sequence AGATCGGAAGAGCACACGTC from the 2nd PCR was removed from 3'-end of reads using cutadapt. Processed reads were aligned to the SecM or EP* reporter plasmid sequences using bowtie version 1.1.2 (*55*). The 5' ends of mapped reads were counted and normalized as reads per million mapped reads (RPM), normalizing for the sequencing depth of each library.

**Mass spectrometric analysis of tmRNA tagging sites**

*Immunoprecipitation and processing of the reporter proteins*

The EP* and the short-SecM (GIRAGP) reporters were expressed in both the wild-type MG1655 strain and the ΔsmrB strain. In addition, a modified tmRNA encoding ANDENYAL**DD** was also expressed from the pAC01 plasmid to stabilize the products of tmRNA tagging. (pAC01 is a derivative of pKW23 (*4*) modified to contain a pBR322 origin). For each of these four samples, reporter protein was purified from three biological replicates as follows: 100 mL LB cultures were grown to $OD_{600}$ = 0.5 and harvested by centrifugation. The pellet was frozen at –80 °C and thawed in 2x CellLytic B cell lysis reagent (Sigma) for 10 minutes. The lysate was clarified by centrifugation for 30 min at 20,000 x g. 50 μL Strep-tactin sepharose beads (IBA) were added to the supernatant and incubated at 4 °C for 1 h. The beads were washed with IP wash buffer (20 mM Tris pH 8.0, 100 mM $NH_4Cl$, 0.4% Triton, 0.1% NP-40) for 5 min at 4 °C four times. Protein was eluted from the beads by shaking at 4 °C in elution buffer (20 mM Tris pH 8.0, 100 mM $NH_4Cl$, 5 mM desthiobiotin) for 1 h. 36 μL of each immunoprecipitated sample was reduced with 1.5 mg/mL DTT in 50 μL of 50 mM tri-ethyl ammonium bicarbonate (TEAB) buffer at 57 °C for 60 min, then alkylated with 10 mg/mL iodoacetomide in 50 μL of 50 mM TEAB buffer in the dark at room temperature for 45 min. The samples were reconstituted in 36 μL of 50 mM HEPES pH 8.5 and digested with 2 ng/μL LysC at 37 °C overnight as described (*62*). Peptides were desalted on Oasis u-HLB plates (Waters), eluted with 60% ACN / 0.1% TFA, dried, and reconstituted with 2% ACN / 0.1% formic acid.

*LC/MS/MS analysis*

Desalted peptides cleaved by LysC were analyzed by liquid chromatography/tandem mass spectrometry (LC/MS/MS). The peptides were separated by reverse-phase chromatography (2% - 90% acetonitrile / 0.1% formic acid gradient over 60 min at 300 nL/min) on an 75 μm x 150 mm ProntoSIL-120-5-C18 H column (Bischoff) using the nano-EasyLC 1200 system (Thermo). Eluting peptides were sprayed into an Orbitrap-Lumos_ETD mass spectrometer through a 1 µm emitter tip (New Objective) at 2.7 kV. Scans were acquired within 360-1700 Da m/z targeting the C-terminal SsrA fusion peptides with no dynamic exclusion. Precursor ions were individually isolated with 0.8 Da (no offset) and fragmented (MS/MS) using HCD activation collision energy 30. Precursor and the fragment ions were analyzed at resolution at 200Da 120,000 AGC target 1xe6, max IT 50ms and 60000, AGC target 1xe5, mx IT118ms, respectively, 3 cycles. Tandem MS/MS spectra were processed by Proteome Discoverer v2.4 (Thermo Fisher) and analyzed with Mascot v.2.6.2 (Matrix Science) using RefSeq2017_83Ecoli and a database with peptides from the nanoLuc-ble reporter protein. Peptide identifications from Mascot searches were processed within the Proteome Discoverer-Percolator to identify peptides with a confidence threshold of a 0.01% False Discovery Rate, based on a concatenated decoy database search to calculate the protein and peptide ratios. Only Peptide Rank 1 were considered.

**Ribosome profiling**

The plasmid encoding the nanoLuc-ble reporter with the short SecM motif (IRAGP) was introduced into four strains: wild-type *E. coli* MG1655, the ΔssrA mutant, the ΔsmrB mutant, and the ΔssrA ΔsmrB double knockout. 200 mL cultures of each strain were grown at 37 °C in MOPS EZ Rich Defined media (Teknova) with ampicillin at 37 °C starting from a 1:100 dilution of an overnight culture to a final $OD_{600}$ = 0.3. The cells were harvested by filtration using a Kontes 99 mm filtration apparatus with a 0.45 µm nitrocellulose filter (Whatman) and then flash frozen in liquid nitrogen. 0.65 mL frozen lysis buffer (20 mM Tris pH 8.0, 10 mM $MgCl_2$, 100 mM $NH_4Cl$, 5 mM $CaCl_2$, 0.1% NP-40, 0.4% TritonX-100, 100 U/mL DNase I (Roche), and 1 mM chloramphenicol) was added to the frozen pellets and The cells were cryogenically pulverized using a Spex 6870 freezer mill with 5 cycles of 1 min grinding at 5 Hz and 1 min cooling. Following lysis, the RNA was digested with MNase and the resulting ribosome footprints were cloned and sequenced as described (*63*).

Custom Python scripts were used to analyze the resulting ribosome profiling data. Raw reads were filtered for quality and trimmed using Skewer v0.2.2. Bowtie v0.12.7 was used to map reads uniquely to genome build NC_000913.2 (allowing two mismatches) and separately to the reporter plasmid sequence after reads mapping to tRNA or rRNA were discarded. Ribosome density was assigned to the 3'-end of reads using read sizes 10–40 nt.

**Polysome profiling**

Cells were cultured at 37 °C in 500 mL of LB (and antibiotics where appropriate) to $OD_{600}$ = 0.5, harvested by filtration using a Kontes 99 mm filtration apparatus with a 0.45 µm nitrocellulose filter (Whatman), and flash frozen in liquid nitrogen. Cells were lysed in lysis buffer (20 mM Tris pH 8.0, 10 mM $MgCl_2$, 100 mM $NH_4Cl$, 5 mM $CaCl_2$, 100 U/mL DNase I, and 1 mM chloramphenicol) using a Spex 6870 freezer mill with 5 cycles of 1 min grinding at 5 Hz and 1 min cooling. Lysates were centrifuged at 20,000 x g for 30 min at 4 °C to pellet cell debris. 10–54% sucrose density gradients were prepared using the Gradient Master 108 (Biocomp) with gradient buffer (20 mM Tris pH 8.0, 10 mM $MgCl_2$, 100 mM $NH_4Cl$, and 2 mM DTT). 5–40 AU of *E. coli* lysate was loaded on top of sucrose gradient and centrifuged in a SW41 rotor at 35,000 rpm for 2.5 h at 4 °C. Fractionation was performed on a Piston Gradient Fractionator (Biocomp). To process each fraction for western blots, proteins were precipitated in 10% trichloroacetic acid (TCA) and the pellets were washed twice by ice-cold acetone, vacuum-dried briefly, resuspended in 5x loading dye, and neutralized with Tris-HCl pH 7.5.

**Purification of *E. coli* SmrB and inactive SmrB mutant ($_{99}$DLH$_{101}$-ALA).**

The plasmids pET24b coding for *E. coli* SmrB and *E.coli* SmrB mutant with N-terminal TwinStrep-tag, TEV cleavage site and FLAG-tag were transformed in *E. coli* stain BL21 (DE3). Cells were grown in 3 L LB medium to mid-log phase ($OD_{600}$ = 0.6) at 37 °C and induced with 1 mM IPTG at 18 °C for 20 h. Cells were harvested by centrifugation at 5,471 *g* and 4 °C for 8 min, resuspended with buffer A (25 mM HEPES/KOH pH 7.8, 300 mM KCl, 5 mM β-Mercaptoethanol, 1:1000 protease inhibitor (pill/mL), 10% glycerol) and lysed using a microfluidizer (Microfluidics M-110L). Cell debris was removed by centrifugation at 30,597 *g* and 4 °C for 20 min. The cleared lysate was then incubated with 5 mL of prewashed Strep-Tactin XT Superflow beads for 1 h. Afterwards the beads were washed with 10 column volumes (CVs) buffer B (25 mM HEPES/KOH pH 7.8, 1 M KCl, 5 mM β-mercaptoethanol, 10% glycerol), with CVs buffer C (25 mM HEPES/KOH pH 7.8, 300 mM KCl, 5 mM β-mercaptoethanol, 10% glycerol), and eluted with 1 CV Buffer D (25 mM HEPES/KOH pH 7.8, 300 mM KCl, 5 mM β-mercaptoethanol, 10% glycerol, 50 mM biotin).

275 µg of TEV protease were added to the elution fraction and incubated on a rotating wheel at 4 °C overnight. To remove the TEV protease, Dynabeads™ (Invitrogen, Thermo Fisher Scientific) were added and incubated for 30 minutes. The cleaved tag was removed by incubation with Strep-Tactin XT Superflow beads. SmrB was concentrated using an Amicon 10k MWCO and subjected to size-exclusion chromatography using a Superdex 75 in buffer C. SmrB-containing fractions were again concentrated and stored at -80 °C.

***E.coli in vitro* translation and isolation of disomes and trisomes**

The VemP-encoding mRNA, which contains the VemP peptide without N-terminal signal sequence, FLAG-tag and a cleavable His-tag, was prepared as described before by PCR amplification, DNA purification, *in vitro* transcription and Phenol/Chloroform precipitation (*34*). RNCs were generated with the PURExpress In Vitro Protein Synthesis Kit (New England Biolabs #E6800S, transcription and translation coupled) using 21 ng of mRNA per 25 µL reaction. 10 reactions were incubated at 30 °C for 35 min and subsequently loaded on sucrose density gradients (25 mM HEPES/KOH pH 7.5, 100 mM KOAc, 10 mM $Mg(OAc)_2$, 0.01% DDM; 10-50% sucrose) and spun in a SW 40 Ti rotor (Beckman Coulter) at 54,322 *g* for 16 h at 4 °C. The gradient was fractionated at a BioComp Gradient Station *ip* using a Triax Flow Cell for UV measurement. The disome and trisomes peak fractions were collected and pelleted by centrifugation in a TLA110 rotor (Beckman Coulter) at 434,513 *g* for 2 h at 4°C. After resuspension in RNC buffer (25 mM HEPES/KOH pH 7.5, 150 mM KOAc, 10 mM $Mg(OAc)_2$, 2 mM DTT), samples were frozen in liquid nitrogen and stored at -80 °C.

***B. subtilis in vitro* translation and isolation of disomes**

The MifM-encoding mRNA, which contains the MifM leader peptide with shortened C-terminus, a defined stalling site, the MifM N-terminal transmembrane segment (TM), a V5-tag and a cleavable His-tag, was prepared as described before by PCR amplification, DNA purification, *in vitro* transcription and Phenol/Chloroform precipitation (*64*). The translation extract was prepared from the *Bacillus subtilis* strain 168 Δhpf ΔssrA ΔSAS1-2 (*65*). Cells were grown in LB medium supplemented with 1% glucose, harvested at an $OD_{600}$ between 0.6 and 0.8 and pelleted by centrifugation at 5,471 *g* and room temperature for 5 min. Afterwards, cells were resuspended in PBS (137 mM NaCl, 2.7 mM KCl, 10 mM $Na_2HPO_4$, 2 mM $KH_4PO_4$, pH 7.4), pelleted again by centrifugation at 5,471 *g* and 4 °C for 15 min and resuspended in as little as possible lysis buffer (10 mM HEPES pH 8.2, 60 mM K glutamate, 14 mM $Mg(OAc)_2$). Cell lysis was performed using a microfluidizer (Microfluidics M-110L) and cell debris was removed by centrifugation at 30,597 *g* and 4 °C for 20 min. The extract was aliquoted and frozen in liquid nitrogen. Activity of the extract as well as Mg buffer concentration was determined using the Luciferase Assay System (Promega).

The *in vitro* translation reaction was performed in 4 x 500 µL reaction volume. 640 µL cell extract were mixed with energy buffer (final concentration in 2 mL: 2% PEG 8000, 50 mM HEPES/KOH pH 8.2, 10 mM $NH_4OAc$, 130 mM KOAc, 30 mM Na-pyruvate, 4 mM Na-oxalate, 50 µg $ml^{-1}$ tRNA (from *E. coli*; Sigma 10109541 001), 0.2 mg ml$^{-1}$ folinic acid, 0.1 µg ml$^{-1}$ creatine kinase, 20 mM creatine phosphate, 4 mM ATP, 3 mM GTP, 0.1 mM amino acid mix, 1 mM DTT, 0.08 U SUPERase•In™ RNase Inhibitor (Invitrogen), 15 mM $Mg(OAc)_2$). After heating the mixture to 32°C for 2 min, 50 µg of mRNA were added to each aliquot and the *in vitro* translation was incubated at 32 °C for 40 min while shaking at 900 rpm. For affinity purification of ribosome nascent chain complexes, the *in vitro* translation was incubated with 400 µL of prewashed TALON metal affinity resin for 45 min on a wheel. The flow-through was collected, beads were washed with 5 CVs buffer A (30 mM HEPES pH7.5/KOH, 250 mM KOAc, 25 mM $Mg(OAc)_2$, 20 mM imidazole, 0.1% DDM) and eluted by incubation for 2 h with 1 CV buffer B (30 mM HEPES pH 7.5/KOH, 250 mM KOAc, 25 mM $Mg(OAc)_2$, 0.1% DDM), 1.1 mg ml$^{-1}$ 3C protease). The sample was loaded on a 10-40% sucrose gradient (30 mM HEPES pH 7.5/KOH, 250 mM KOAc, 25 mM $Mg(OAc)_2$, 0.1% DDM, 10-40% (w/v) sucrose) and spun in a SW 40 Ti rotor (Beckman Coulter) at 54,322 *g* for 16 h at 4°C. The disome peak fractions were combined pelleted by centrifugation in a TLA110 rotor (Beckman Coulter) at 434,513 *g* for 2 h at 4°C. The pellet was resuspended in buffer C (25 mM HEPES pH 7.5/KOH, 150 mM KOAc, 10 mM $Mg(OAc)_2$, 2 mM DTT), frozen in liquid nitrogen and stored at -80 °C.

**Nuclease Assay**

Purified *E. coli* disomes were mixed with 10 times molar access of protein (SmrB wt/ SmrB mut). As a control the same volume of buffer C (25 mM HEPES pH 7.5/KOH, 150 mM KOAc, 10 mM $Mg(OAc)_2$, 2 mM DTT) was added to disomes. Samples were incubated at 30°C for 3 h and then loaded on sucrose density gradients (25 mM pH 7.5 HEPES-KOH, 100 mM KOAc, 10 mM Mg(OAc)2, 0.01% DDM; 10-50% sucrose). The gradients were spun in a SW 40 Ti rotor (Beckman Coulter) at 54,322 *g* for 16 h at 4°C. The gradient was fractionated at a BioComp Gradient Station *ip* using a Triax Flow Cell for UV measurement.

**Cryo-EM analysis**

*Data collection and processing of the E. coli disome and E. coli trisome.*

A volume of 3.5 µL was applied to 2 nm pre-coated Quantifoil R3/3 holey carbon support grids and vitrified in liquid ethane using a Vitrobot mark IV (FEI Company, Netherlands) (wait time 45 s, blotting time 2 s). 2,437 and 14'849 movies were collected for the *E. coli* disome and trisome sample, respectively. Data were collected on a Titan Krios TEM using a Falcon II DED at 300 kV, with an electron dose of 2.5 $e^-/Å^2$ per frame for 16 frames (defocus range of 0.5 to 4 µm). The magnified pixel size was 1.09 Å/pixel. For the *E coli* disome sample, frames were gain corrected, aligned and summed using MotionCor2 (*66*) and CTF parameters were determined using CTFFIND (*67*). Particles were picked using Gautomatch (http://www.mrc-lmb.cam.ac.uk/kzhang/). The particles were extracted and processed following the standard workflow in RELION 3.1 (*68*). The particles containing the stalled ribosome of the disomes were extracted with a box size of 380 pixel, imported to Cryosparc v3.2.0 (*69*) and refined to a final resolution of 3.8 Å (Fig S5). For the *E. coli* trisome, data were processed following the standard workflow in cryoSPARC v3.2.0 (*69*).

*Data collection and processing of the B. subtilis disome*

All samples were vitrified as described above. Two datasets of 8,842 and 19'354 movies were collected on a Titan Krios TEM using a Falcon II DED at 300 kV, with an electron dose of 2.5 $e^-/Å^2$ per frame for 16 frames (defocus range of 0.5 to 4 µm). The magnified pixel size was 1.084 Å/pixel. All frames were gain corrected and subsequently aligned and summed using MotionCor2 (*66*). The data were processed following the standard workflow in cryoSPARC v3.2.0 (*69*).

*Sample preparation, data collection and processing of the E. coli disome SmrB complex.*

For reconstitution, disomes and SmrB mutant ($_{99}DLH_{101}$-ALA) were thawed on ice. Disomes were mixed with

10 times molar access of protein, incubated for 10 min at room temperature and subsequently analysed by cryo-EM8,350 movies were collected on a Titan Krios at 300 kV recorded on a K2 Summit direct electron detector (DED) with an electron dose of approx. 1.06 $e^-/Å^2$ per frame for 40 frames (defocus range of 0.5 to 3.5 µm). The magnified pixel size was 1.059 Å/pixel. All frames were gain corrected and subsequently aligned and summed using MotionCor2 (*66*). The data were processed following the standard workflow in cryoSPARC v3.2.0 (*69*). The processing scheme and final local resolution for SmrB are shown in Fig S7.

**Model building and refinement**

The *E. coli* disome model was prepared by rigid body docking of the model from Su *et al.* (PDB code 5NWY (*34*)). uL9 was taken from PDB-6WD1 (*70*) and the N-terminal part (residues 1-52) and the C-terminal part (residues 53-149) were rigid body docked individually and rejoined to match the bridged conformation of uL9 in the stalled ribosome. uL1 and the L1-stalk rRNA (nucleotides 2099-2190 were taken from PDB-6WD1 and rigid body docked into the cryo-EM density. tRNA-Phe was taken from PDB-3L0U (*71*) and rigid body docked into the A-sites of stalled and collided ribosomes although the identity of the tRNA in the A-site of the collided could not be determined. uS1 was taken from PDB-6BU8 and docked into the density map of the collided ribosome. The resolution of the mRNA between the two ribosomes was insufficient for modeling with nucleotide precision. Despite that, the mRNA of PDB-5NWY was extended at the 5'-end according to the sequence of the construct to provide an approximation of the number of nucleotides stretching from the exit of the stalled to the entry of the collided ribosome. The SmrB model was prepared using Alphafold 2 (AF2) (*39*) predictions and Mmseqs2 (*72*) for multiple sequence alignment of SmrB alone and of SmrB fused to the ribosomal interaction partner uS2 as described in Fig S8. AF2 predictions were performed using an API provided by the Söding lab.

The *E. coli* trisome model was prepared by docking the two copies of the stalled 70S from the disome in the first and second ribosome and one copy of the collided 70S in the third ribosome. The *B. subtilis* disome model was prepared by docking two copies of the MifM-stalled ribosome complex (PDB 3J9W; (*64*)) into the cryo-EM map.

All model adjustments were performed using coot (*73*). Structural figures were prepared using ChimeraX (*74*).

**Figure S1**

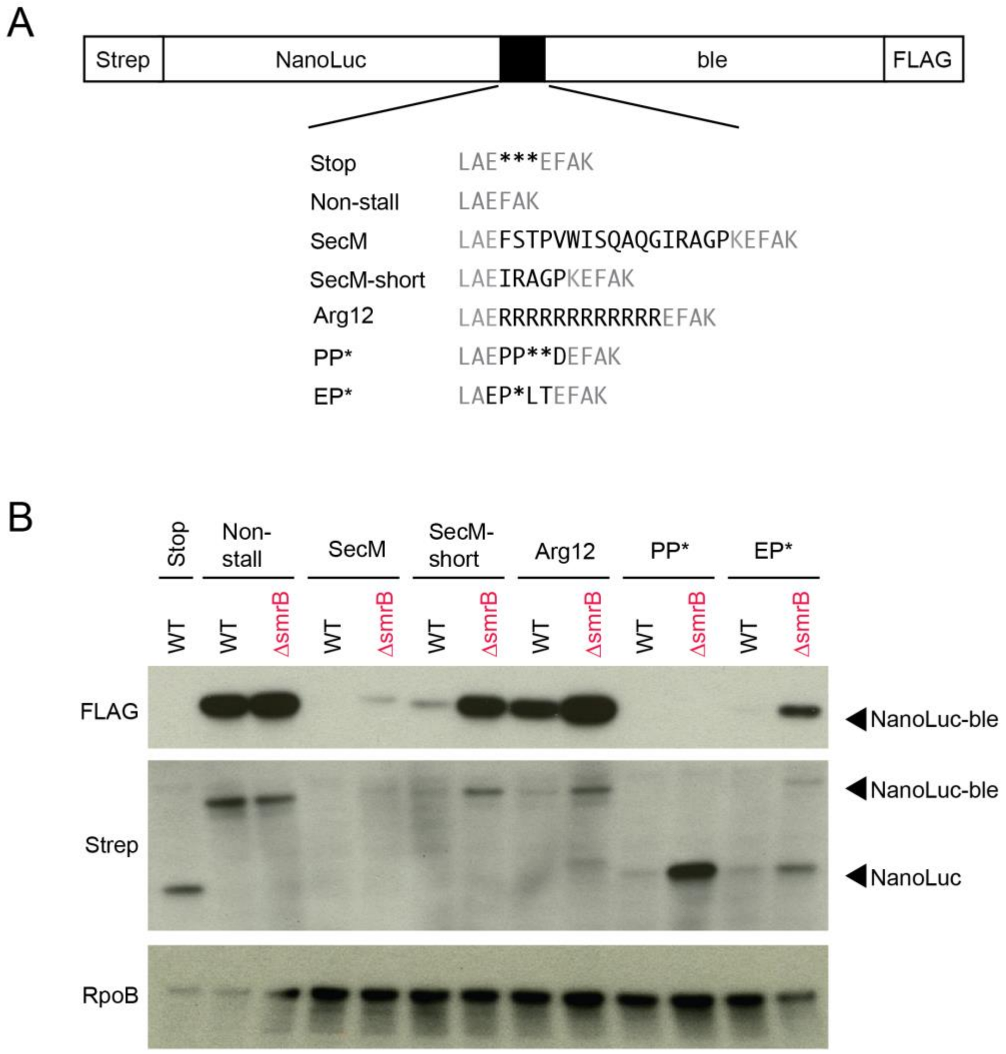


**Supplemental Figure 1. SmrB is a general quality control factor**

**(A**) Additional reporters to study ribosome rescue in *E. coli* with various stall motifs. (**B**) The expression of full-length NanoLuc-Ble protein was monitored an anti-FLAG antibody; anti-Strep antibodies reveal both full-length NanoLuc-Ble and truncated NanoLuc proteins. The RpoB protein serves as a loading control.

**Figure S2**

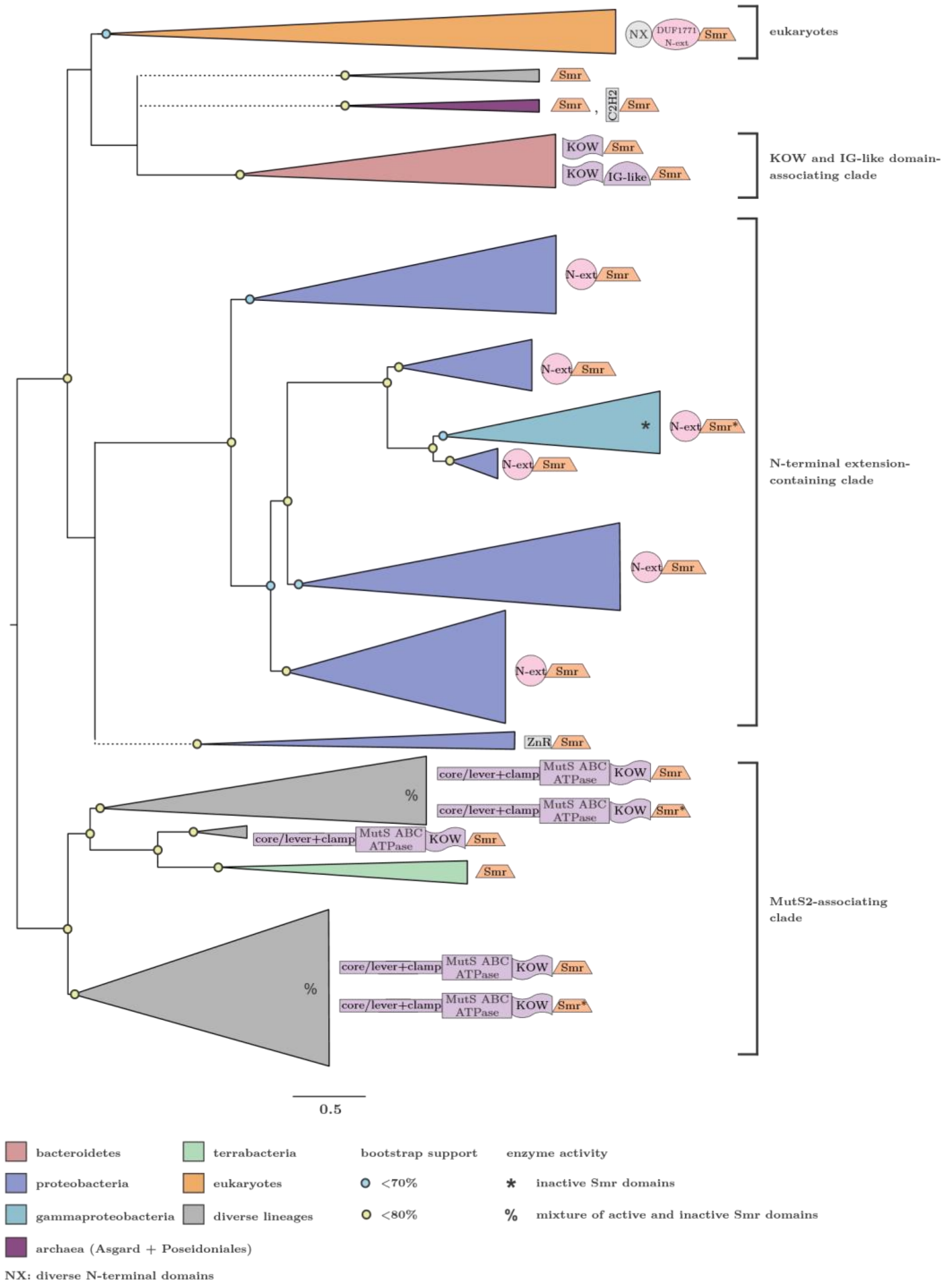


**Supplemental Figure 2. Phylogenetic tree of SMR domain proteins**

Stylized phylogenetic tree depicting relationships between SMR domain clades. Clades with indicated bootstrap support are marked with circles. Clade names are given to the right of the tree. Dotted lines indicate positions with little or no bootstrap support.

**Figure S3**

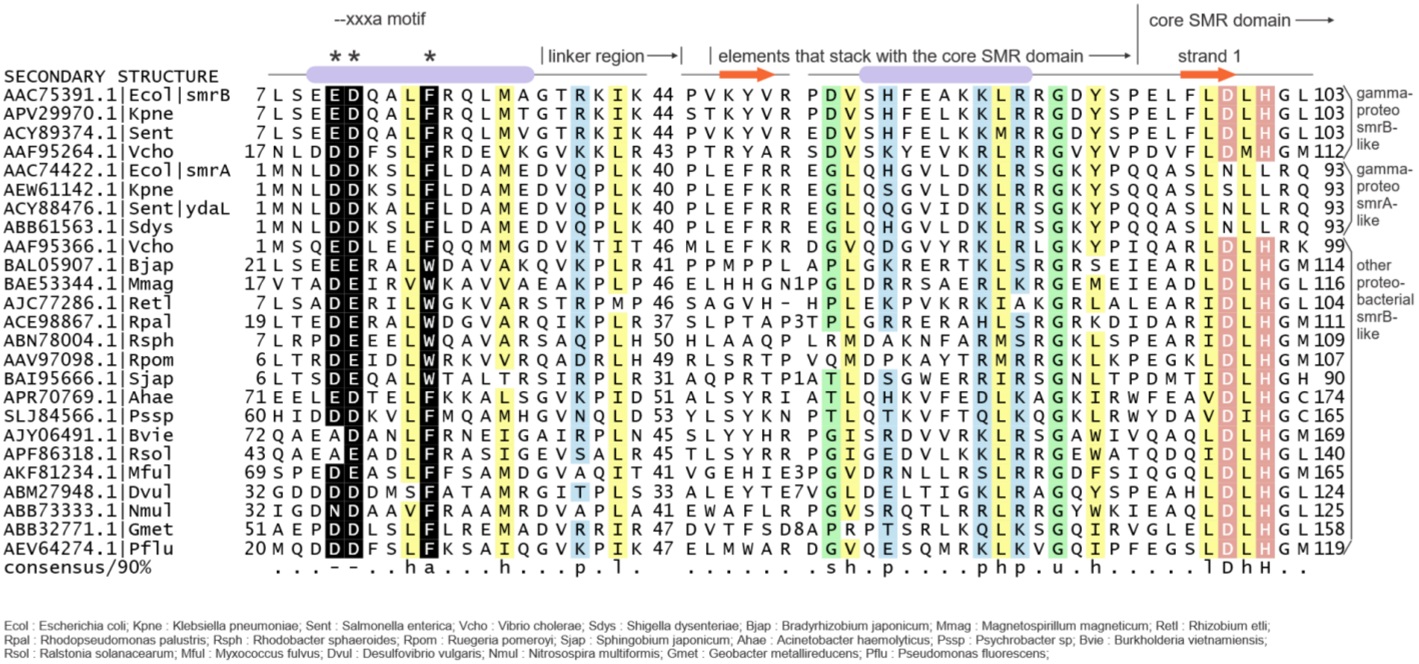


**Supplemental Figure 3. Multiple alignment of the conserved regions in the N-terminal extension of SMR proteins from proteobacteria.** Columns in the alignment are shaded and labeled according to biochemical character: -, negatively charged; h, hydrophobic in yellow; a, aromatic; p, polar in blue; l, aliphatic in yellow; s, small in green; u, tiny in green. Residue positions in the --xxxa motif are colored in white and shaded in black, marked by asterisks above the alignment. Residue positions forming part of the active site of the core SMR domain are colored in white and shaded in red. Sequences are labeled with NCBI accession number and organism abbreviation, abbreviations are provided below alignment. Secondary structure provided at top of alignment. Numbers to left and right of alignment denote positioning of the region. Internal numbers give the size of excised variable insert regions.

**Figure S4.**

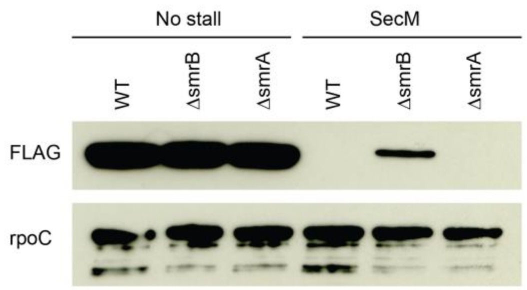


**Supplemental Figure 4. Loss of SmrA does not affect expression of the stalling reporter**
The level of SecM reporter protein was monitored using an anti-FLAG antibody. The RpoC protein serves as a loading control.

**Figure S5.**

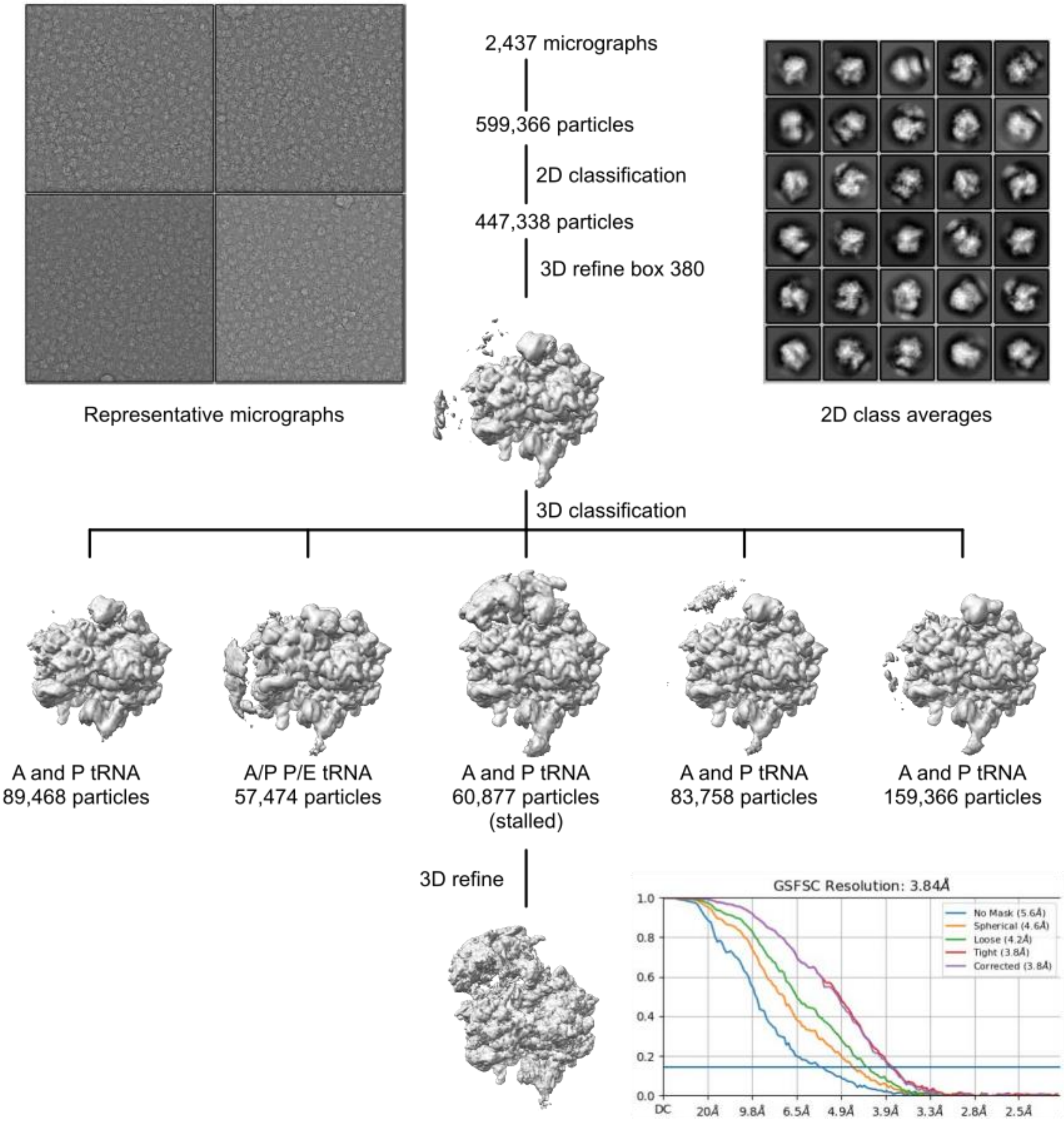


**Supplemental Figure 5. Cryo-EM data processing for the *E. coli* disome sample.** Shown are the classification scheme, representative micrographs, 2D class averages and the Gold standard Fourier Shell Correlation (GSFSC) curve for the final volume containing the 70S stalled ribosome as well as the 30S of the collided ribosome.

**Figure S6**

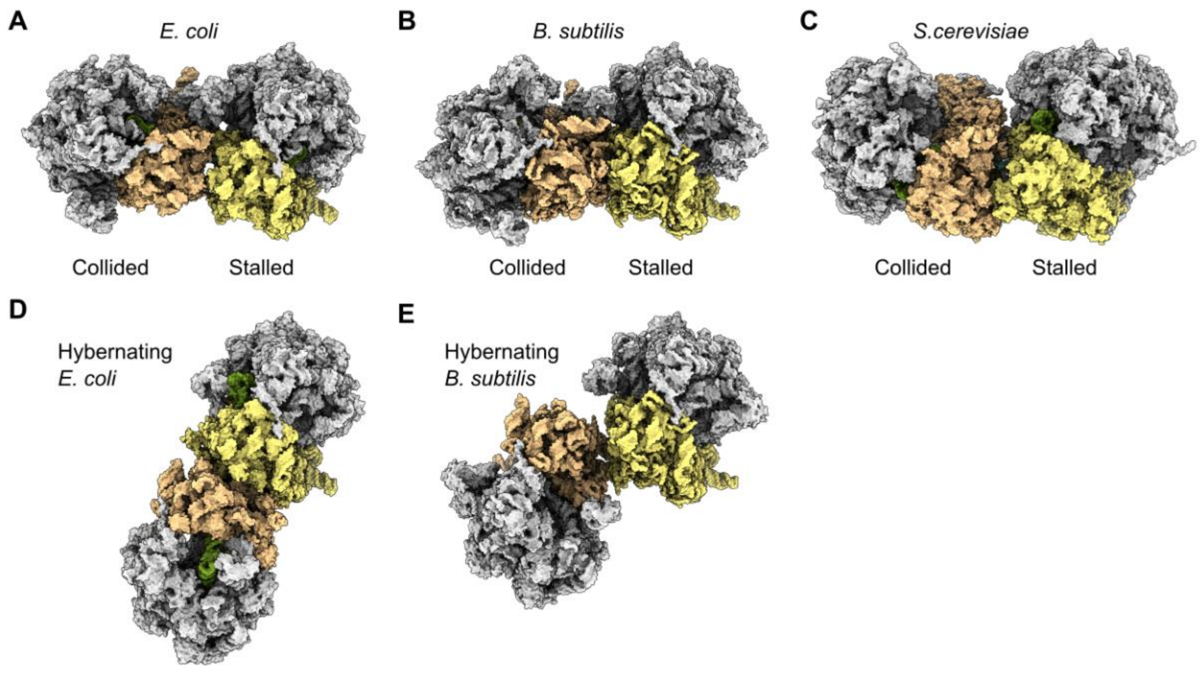


**Supplemental Figure 6. Comparison of different disomes.** The *E. coli* and *B. subtilis* disomes from ribosome collision are compared to the S. cerevisiae ribosome collision disome and hibernation disomes.

**Figure S7**

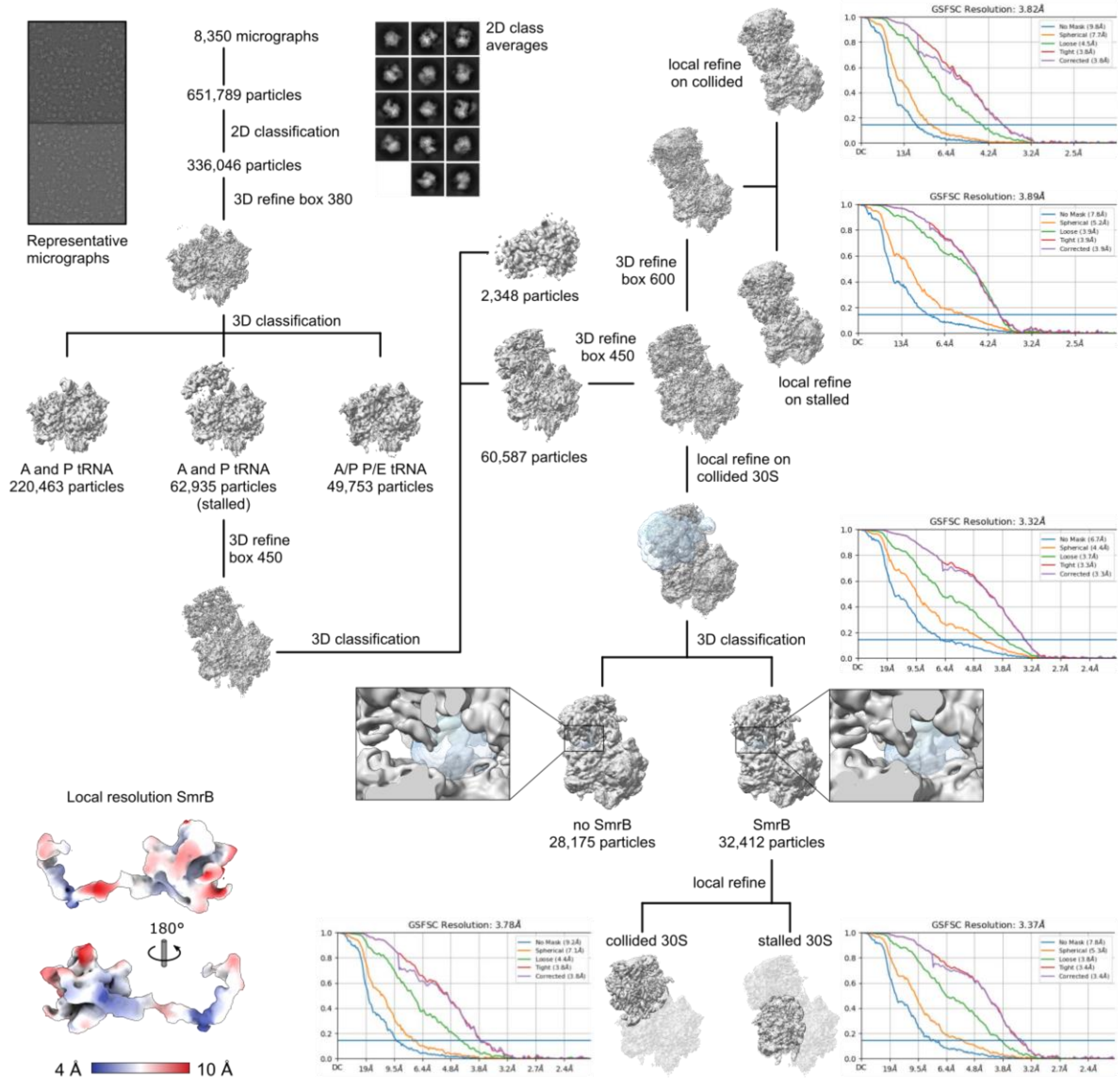


**Supplemental Figure 7. Cryo-EM data processing for the *E. coli* disome sample.** Shown are the classification scheme, representative micrographs, 2D class averages and the Gold standard Fourier Shell Correlation (GSFSC) curve for the respective 3D reconstructions. The segmented density for SmrB is colored according to local resolution.

#### Figure S8

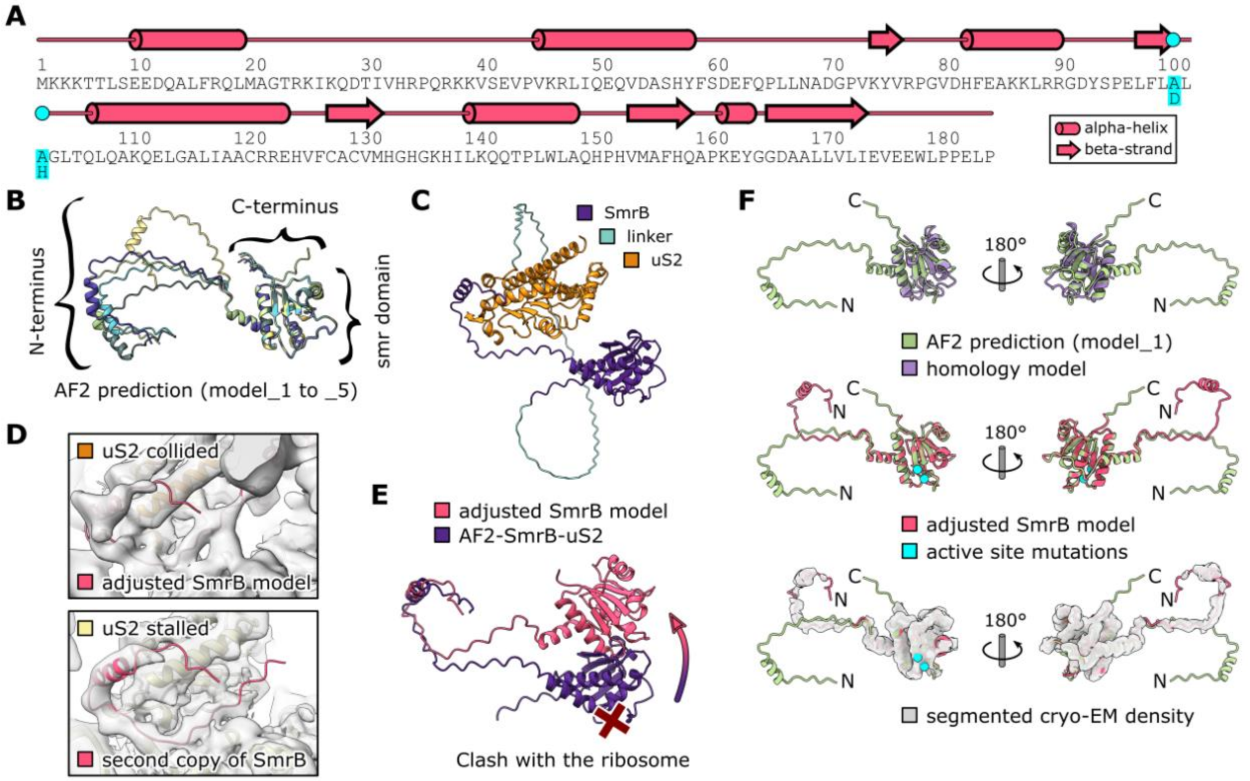


**Supplemental Figure 8. Structural model of SmrB.** (**A**) Secondary structure of SmrB. The DLH to ALA mutation is indicated. (**B**) AF2 prediction models 1-5. The SMR domain is predicted with high confidence, while the linker to the N-terminal helix appears flexible. (**C**) AF2 prediction of the interaction between SmrB and uS2. For this prediction uS2 was fused to the C-terminus of SmrB with a glycine serine linker (39 copies of GS). The prediction shows the N-terminal helix of SmrB folded back onto uS2. (**D**) Top: Cryo-EM density and adjusted model of the SmrB. Bottom: Cryo-EM density and rigid body docked model of the N-terminus of SmrB from the collided 30S onto the stalled 30S. A second copy of SmrB was found anchored to uS2 of the stalled ribosome. However, there was no density for the SMR domain of the second SmrB, indicating a high degree of flexibility due to the lack of another ribosome in front of the stalled one. (**E**) Adjustment of the AF2 predicted model of SmrB-uS2. Without adjustment according to the cryo-EM density (as shown in D) the SMR domain would clash with the ribosome. (**F**) Comparison of the AF2 prediction, the homology model and the adjusted model of SmrB. Compared to the AF2 prediction, the homology model is missing the two N-terminal helices and most of the loops are slightly different (top). The AF2 prediction almost perfectly matched the cryo-EM density map and the corresponding adjusted model (middle and bottom). Only the catalytic loop (carrying the active site mutations) had to be slightly adjusted to prevent clashes with the mRNA. The N-terminus was adjusted as discussed above.

**Figure S9**

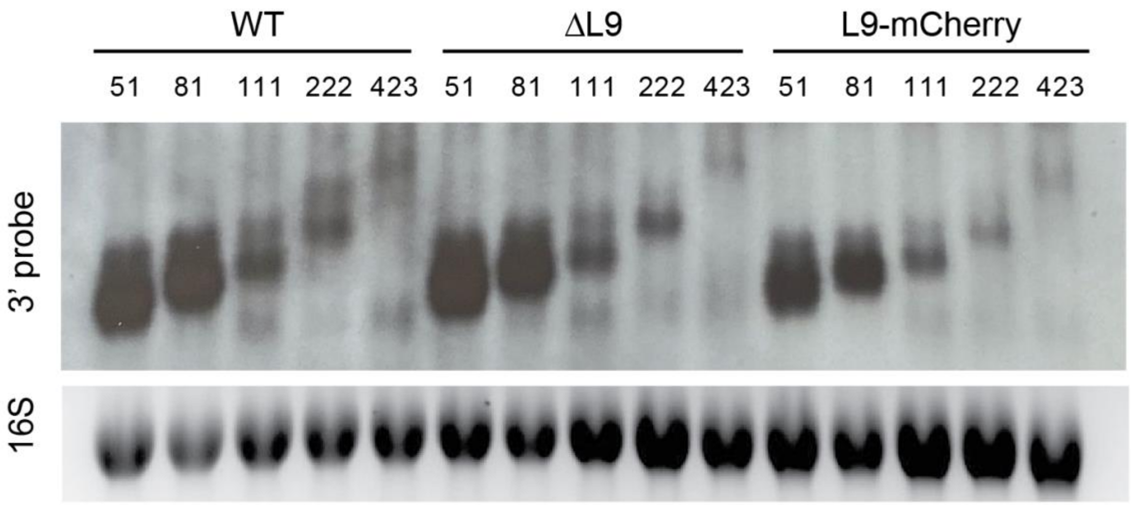


**Supplemental Figure 9. RNA processing via collision is independent of the L9 bridge**
Northern blots using the 3'-probe against the CRP reporters with the short SecM stalling motif in wild-type cells, L9-deletion strain (ΔL9), and a strain where mCherry is fused to the C-terminus of L9 (L9-mCherry). Ethidium bromide staining of 16S rRNA serves as a loading control.
